# Multi-source DOM inputs dictate net extra carbon mineralization in eutrophic lakes via non-linear co-metabolism effects

**DOI:** 10.64898/2026.11.10.760625

**Authors:** Jiaxing Tian, Jingmin Wang, Yujia Chen, Wenxin Wei, Xudong Dai, Ziqian Gong, Qiuyuan Ren, Chao Wu, Yaoqing Peng, Jicheng Zhong, Fei Yang, Xiaoguang Xu, Jianjun Wang

**Affiliations:** State Key Laboratory of Lake Science and Environment, Nanjing Institute of Geography and Limnology, Chinese Academy of Sciences, Nanjing, 211135, China; School of Environment, Nanjing Normal University, Nanjing 210023, China; Ministry of Ecology and Environment, Nanjing Institute of Environment Sciences, Nanjing, 210042, China

**Author notes:** Corresponding authors. 299, Chuangzhan Road, Qilin Street, Jiangning District, Nanjing, 211135, China Corresponding author for editorial correspondence: Xiaoguang Xu.

**Keywords:** Dissolved organic matter, eutrophic lake, carbon mineralization, greenhouse gas, co-metabolism

## Abstract

Microbially mediated decomposition of dissolved organic matter (DOM) is a key process in the carbon cycle of inland lakes. Exogenous DOM inputs can alter the decomposition of resident lake-water DOM through co-metabolic effects (CE), yet how DOM source and addition level jointly regulate this process remains poorly understood, especially in eutrophic lakes. Here, we selected Lake Taihu, a representative hypereutrophic lake, and conducted incubation experiments by separately adding DOM derived from rainwater, leaves of aquatic plants, terrestrial litter, and treated sewage effluent to algal-rich lake water at matched carbon-addition levels. Water chemistry and DOM optical properties were used to identify factors associated with source- and concentration-dependent CE. DOM from aquatic plants and rainwater, which exhibited relatively strong aromatic and humified characteristics, induced positive CE. In contrast, terrestrial litter-derived DOM produced a weaker positive CE, while sewage-derived DOM, characterized by stronger fresh autochthonous and microbial signatures and lower average molecular weight, induced negative CE. The magnitude of CE varied nonlinearly with addition level, peaking at intermediate concentrations. CE was closely related to DOM aromaticity, humification, fresh autochthonous characteristics, and nitrogen conditions. A source-resolved scenario analysis combining estimated DOM-C inputs with source-specific CE indicated that the negative contribution of sewage-derived DOM could offset much of the positive contribution from natural sources, although the combined effect remained a potential net increase in carbon mineralization. These findings demonstrate that DOM from different environmental sources can induce CE in opposing directions in bloom-affected Lake Taihu water and that CE magnitude depends nonlinearly on loading level.

## 1. Introduction

Although inland waters cover only approximately 3.7% of the global ice-free land surface, they are active interfaces for the transformation, burial, and release of terrestrial organic carbon [1, 2]. Dissolved organic matter (DOM), a central carrier of carbon in lakes, is highly heterogeneous in composition, reactivity, and microbial bioavailability [3, 4, 5]. In eutrophic lakes, water-column mixing and sediment-water exchange continually bring algal DOM into contact with exogenous DOM delivered by rainfall, littoral vegetation, and anthropogenic discharges. These inputs can alter microbial utilization of lake-water DOM and connect otherwise distinct carbon pools through co-metabolic effects [6, 7]. Based on long-term observations from 2000 to 2022, annual CO₂ emissions from Lake Taihu are estimated at 1.1 ± 0.3 Tg, with the highest emissions primarily occurring in the northwestern region of the lake, where riverine DOM inputs are influential [8]. Resolving how exogenous DOM inputs alter DOM metabolism is therefore important for understanding organic-carbon fate and the role of lakes as carbon sources or sinks.

The co-metabolic effect (CE) refers to the process whereby microorganisms using newly supplied organic carbon simultaneously alter the decomposition and transformation of organic carbon already present in an ecosystem [6, 7]. CE is evaluated by comparing the observed changes in organic carbon in a mixed system with the theoretical changes expected under a no-interaction assumption [9, 10, 11]. A positive CE indicates enhanced decomposition of the resident organic carbon pool, whereas a negative CE indicates suppression. Aquatic studies have reported positive, negative, weak, and undetectable CEs [9, 11, 12], highlighting the strong context dependence of this process.

Exogenous DOM may enhance the decomposition of resident DOM by supplying energy and stimulating extracellular enzyme activity. However, it may also inhibit decomposition when microorganisms preferentially utilize the newly introduced substrate. The interplay between these opposing processes ultimately dictates the direction and magnitude of CE [6, 13, 14]. Most aquatic CE studies have used glucose, acetate, or a single algal substrate; however, direct cross-study comparisons are limited by differences in recipient DOM, substrate type, addition levels, and CE calculations [9, 12, 14]. CEs have also been observed for DOM from real environmental sources, including rainwater, plant leachates, litter, and sewage [15, 16, 17]. However, direct evidence remains scarce regarding whether equal quantities of DOM-C from different sources yield equivalent co-metabolic effects in bloom-affected eutrophic waters and how the CE from each source changes with input concentration. Large, shallow eutrophic lakes receive episodic rainfall inputs, seasonal releases from plants and litter, and continuous inputs of treated sewage effluent [18, 19, 20], providing an ideal environment to investigate how DOM source and loading jointly regulate carbon decomposition in mixed systems.

In this study, DOM derived from typical sources, including rainwater, leaves of the aquatic plant *Potamogeton malaianus*, terrestrial litter of Chinese tallow tree (*Triadica sebifera*), and treated sewage effluent, was separately added to algal-rich water from eutrophic Lake Taihu at matched carbon-addition levels. CE was quantified using a mass-balance approach, and the results were integrated with DOM optical properties, water chemistry, and greenhouse gas measurements to identify factors associated with source- and concentration-dependent effects. Source-specific CE estimates were then combined with estimated DOM-C inputs to assess their potential contributions to carbon mineralization in Lake Taihu. This study addressed two primary questions: (1) Do different environmentally derived DOM sources generate different CEs in the same bloom-affected water? and (2) How does the CE of each source vary with the level of exogenous DOM input?

## 2. Materials and methods

### 2.1 Sample collection and DOM preparation

Surface water collected from a cyanobacterial bloom accumulation zone in Lake Taihu, China, was used as the recipient lake water. Four types of exogenous DOM were prepared from rainwater, leaves of the aquatic plant *Potamogeton malaianus*, terrestrial litter of Chinese tallow tree (*Triadica sebifera*), and final effluent from a sewage treatment plant. Both the recipient lake water and rainwater were collected at the Yixing Cyanobacterial Bloom Observation Station (120°00′42″ E, 31.4352° N). Leaves of *P. malaianus* and terrestrial litter of T. sebifera were collected from the adjacent littoral zone, while treated effluent was obtained from the final discharge outlet of a municipal sewage treatment plant in Yixing City. To capture wet-deposition DOM entering the lake surface directly, rainwater was sampled during a continuous rainfall event from 28 to 30 July 2025, following an event-based protocol [15]. The collector was placed in an open, unobstructed area away from canopy drip, roof runoff, and road splash. Total rainfall was 48.3 mm. Four subsamples were collected at approximately 6 h intervals and combined to represent rainwater-derived DOM. Rainwater was collected using an acid-washed stainless-steel collector that had been thoroughly rinsed with Milli-Q water. Leaves of *P. malaianus* and terrestrial litter of T. sebifera were used to represent DOM derived from typical littoral aquatic vegetation and surrounding terrestrial litter, respectively. After sediment and surface debris were removed, the plant materials were cut into pieces approximately 1 cm in length. Following previously published extraction methods [15, 16], the aquatic plant leaves and terrestrial litter were mixed with Milli-Q water at solid-to-liquid ratios of 1:10 and 1:5 (g dry weight: mL), respectively. Dry-weight equivalents were used for all solid masses, with the fresh weight of *P. malaianus* leaves converted accordingly. Samples were extracted in the dark at 25 °C for 24 h and then centrifuged at 10,000 rpm for 30 min. The source-specific solid-to-liquid ratios were selected based on established procedures to ensure sufficient stock DOC concentrations for the subsequent carbon-addition gradient. Sewage-derived DOM consisted of final effluent collected directly from the treatment-plant discharge outlet.

Rainwater, aquatic plant extract, terrestrial-litter extract, treated effluent, and recipient lake water were sequentially filtered through precombusted GF/F filters and 0.22-μm polycarbonate membranes, transferred to precleaned amber-glass bottles, and stored at 4 °C in the dark. A microbial inoculum was prepared from the same batch of fresh Lake Taihu water by membrane retention followed by shaking and resuspension [21]. Specifically, lake water was filtered through a 0.22-μm polyethersulfone membrane, and the retained microorganisms were collected. The membrane was then placed in 30 mL of the corresponding lake-water filtrate and shaken at 150 rpm for 30 min to resuspend the microorganisms. The resulting suspension was passed through a 1.6-μm glass-fiber filter to remove larger particles and planktonic consumers, and the filtrate was used as the inoculum. All incubation systems received 10% (v/v) of the same inoculum to maintain a common microbial source, and the DOC introduced with the inoculum was included in the measured day-0 concentration of each system.

The extraction ratios of aquatic plants and terrestrial litter were 10 mL and 5 mL of Milli-Q water per gram of dry-weight equivalent, respectively. After centrifugation, the supernatants were collected as aqueous leachates and subsequently subjected to the sequential filtration process described above; intact plant fragments and litter residues were excluded from the incubations. Rainwater and final effluent were filtered and then used directly, without solid–liquid extraction. These four filtered aqueous preparations constituted the source-DOM stock endmembers. DOC concentrations in these stocks were measured prior to calculating the volumes required for the carbon-addition treatments [15].

### 2.2 Incubation experiment

The incubation experiment comprised four DOM sources and five addition levels. Prior to incubation, DOC concentrations in the source stocks were measured, and the volume of each stock required to achieve final exogenous-carbon additions of 4, 8, 16, 32, or 48 μM C was calculated accordingly. For each source and addition level, the mixed treatment consisted of exogenous DOM and recipient lake water. A lake-water endmember control and four source-DOM endmember controls were established to quantify DOC changes when lake water and each source stock were incubated separately. All endmember controls received the same proportional volume of microbial inoculum. Mixed treatments were adjusted to the same final volume with Milli-Q water. DOC changes observed in the lake-water and source-DOM endmembers were used to calculate theoretical DOC removal under the no-interaction assumption [12, 15, 16].

Each source-DOM endmember control was prepared by combining 225 mL of the corresponding filtered source stock with 25 mL of a common microbial inoculum, yielding a total liquid volume of 250 mL. These controls represented the source-stock endmembers, rather than source DOM diluted to each mixed-treatment addition level. They were incubated under the same conditions and sampled on the same schedule as the mixed treatments. The measured day-0 DOC concentrations of these controls included the carbon introduced with the inoculum.

Incubations were conducted in sealed 400-mL glass bottles containing 250 mL of liquid and a 150-mL headspace. Each treatment had three independent replicates, received 10% (v/v) of the same inoculum, and was incubated in the dark at 25 °C for 32 d. On days 0, 1, 7, 17, and 32, 10 mL of headspace gas and 20 mL of incubation liquid were collected from each bottle. CE was calculated at each sampling time, with day 32 used as the primary endpoint.

Following the mass-balance approach used in previous mix-incubation studies of aquatic DOM [15, 16], CE was calculated as the deviation between the measured DOC removal in a mixed system and the theoretical DOC removal expected in the absence of interactions. For each DOM source, addition level, and sampling time, theoretical DOC removal was calculated by weighting DOC removal in the lake-water control and the corresponding source control by their volumetric contributions to the mixed system:

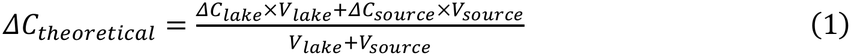

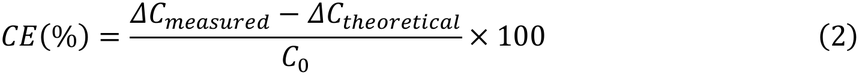

where ΔC_measured_ is the measured DOC removal from day 0 to the corresponding sampling day in the mixed system; ΔC_lake_ and ΔC_source_ are DOC removals over the same period in the lake-water and source controls, respectively; V_lake_ and V_source_ are the volumes of lake water and exogenous-DOM stock in the mixed system, respectively; and C_0_ is the measured DOC concentration in the mixed system on day 0. CE > 0 indicates that DOC removal in the mixed system exceeded the no-interaction expectation, whereas CE < 0 indicates that DOC removal was lower than expected. Equation (1) is a carbon-amount balance expressed in concentration units. Multiplying each endmember DOC concentration decrease by its contributing volume gives the corresponding amount of carbon removed; summing the ΔClake × Vlake and ΔCsource × Vsource terms and dividing by Vlake + Vsource converts that amount back to a theoretical concentration decrease. Unequal DOC concentrations in lake water and source stocks are therefore not ignored. An equivalent mass-based calculation would multiply each endmember’s initial carbon amount by its fractional DOC loss, yielding the same concentration-by-volume terms. Applying an additional initial-DOC mass fraction to the absolute concentration decreases would instead weight concentration twice. This two-endmember formulation follows Zhuang et al. [15]; concentration changes and volume weights must refer to a consistent dilution and inoculum basis.

### 2.3 Physicochemical analyses

Concentrations of CO_2_, CH_4_, and N_2_O were measured using a gas chromatograph (Agilent 7890B, Agilent Technologies, USA). Gas concentrations were converted to fluxes based on the incubation temperature and actual headspace volume. Changes in gas concentration between adjacent sampling times were corrected for concurrent changes in the lake-water control and converted using the corresponding headspace volume. Cumulative values on day 32 were calculated as the sum of net changes across incubation intervals. pH and dissolved oxygen (DO) were measured with a multiparameter water-quality meter (YSI ProDSS, Xylem, USA). The pH probe was calibrated with standard buffers at pH 4.01, 7.00, and 9.18, and the DO probe was calibrated at 100% saturation in water-saturated air.

Incubation liquid was filtered through GF/F filters for water-chemistry, DOC, and DOM optical analyses. Permanganate chemical oxygen demand (COD_Mn_) was measured by acidic potassium-permanganate oxidation. Total nitrogen (TN) was measured by alkaline persulfate digestion followed by ultraviolet spectrophotometry. Total phosphorus (TP) and orthophosphate phosphorus (PO_4_^3-^-P) were measured by persulfate digestion followed by ammonium-molybdate spectrophotometry and by ammonium-molybdate spectrophotometry, respectively. Ammonium nitrogen (NH_4_⁺-N), nitrite nitrogen (NO_2_^-^-N), and nitrate nitrogen (NO_3_^-^-N) were measured by salicylate spectrophotometry, N-(1-naphthyl) ethylenediamine spectrophotometry, and ultraviolet spectrophotometry, respectively (HJ/T 346-2007) [22]. Filtrate for DOC analysis was acidified to pH < 2 with HCl and analyzed by high-temperature catalytic oxidation using a total organic carbon analyzer (TOC-L CPH, Shimadzu, Japan). Each sample underwent two or three automated measurements and was reanalyzed when the coefficient of variation exceeded 5%.

The DOC quality assurance and quality control protocol addressed sample handling and analytical repeatability. Prepared source stocks were stored in precleaned amber-glass bottles at 4 °C in the dark, as described in Section 2.1. Aliquots for DOC analysis were filtered and acidified as described above. Analytical precision was evaluated from the two or three automated measurements of each sample using the coefficient of variation (100 × standard deviation/mean); samples were reanalyzed when this value exceeded 5%. These instrumental repeats were distinct from the three independent incubation replicates. Control of handling contamination, preservation, and analytical repeatability is important for reliable DOC measurements [23].

DOM absorbance spectra were measured from 200 to 800 nm using a UV-visible spectrophotometer (UV-2600, Shimadzu, Japan), with Milli-Q water as the blank. Absorbance was corrected against the 700–800 nm baseline and converted to the Napierian absorption coefficient a(λ). The indices a_254_, a_350_, E2/E3, S_275-295_, S_R_, and SUVA_254_ were then calculated [24, 25].

Three-dimensional excitation-emission matrices (EEMs) were measured using a fluorescence spectrophotometer (Cary Eclipse, Agilent Technologies, USA) over excitation wavelengths of 200–450 nm and emission wavelengths of 250–500 nm. EEMs were corrected sequentially by subtraction of the Milli-Q water blank, Raman normalization, inner-filter-effect correction, and removal of scattering regions. The corrected spectra were used to calculate the fluorescence index (FI), humification index (HIX), biological index (BIX), and β:α ratio, which describe DOM source, degree of humification, and the contribution of fresh autochthonous material [26, 27, 28, 29]. Parallel-factor analysis (PARAFAC) was subsequently used to resolve fluorescent components. Model stability was evaluated using component number, residuals, and split-half analysis [30, 31]. Model validation and component interpretation are provided in Supplementary Method S1, Fig. S3, and Table S4.

### 2.4 Data analysis

One-way analysis of variance (ANOVA) was used to compare CE, absorbance and fluorescence indices, water-chemistry variables, and gas variables among the four exogenous-DOM treatments. Principal-component analysis (PCA) was performed after z-score standardization of absorbance and fluorescence indices [32]. The principal components were interpreted based on the direction and magnitude of variable loadings, and the axis dominated by aromaticity and humification indices was interpreted as a composite DOM aromaticity-humification gradient. To characterize changes in CE along the ordered addition gradient, polynomial regressions containing linear and quadratic concentration terms were fitted separately for each DOM source. Model residuals and goodness of fit were used to evaluate the relationships. Details of the PCA, regression models, and diagnostics are provided in Supplementary Method S3. These regressions were used to describe response trends within the experimental concentration range. Associations between CE and water chemistry, DOM optical indices, and gas variables were assessed using Spearman rank correlations. Random forest regression was used to identify the principal variables explaining day-32 CE [33]. Candidate predictors included the added-DOC concentration, DOM absorbance and fluorescence indices, PARAFAC components, pH, DO, and major nutrient variables. Cumulative gas production was treated as a carbon-mineralization response parallel to CE and was therefore excluded from the predictor set. Predictor contributions were expressed as permutation importance, and model performance was evaluated using cross-validated R^2^ and root-mean-square error (RMSE). All statistical analyses were performed in R 4.3.2 using two-sided tests and a significance threshold of *p* < 0.05. Data screening, multiple comparisons, random forest parameters, and model validation are detailed in Supplementary Method S3.

Annual inputs from the four DOM sources were estimated using a source-resolved mass-balance approach. Rainwater-derived DOM-C input was calculated from the surface area of Lake Taihu, annual precipitation, and a literature-derived rainwater DOC coefficient [18]. Sewage-derived DOM-C input was calculated from treated effluent discharge and a literature-derived effluent DOC coefficient [34, 35]. Aquatic plant-derived DOM-C input was calculated from aquatic-vegetation area, in situ DOC release rates, and effective daylight hours during the growing season [20, 36]. Potential production of terrestrial-litter DOC was first estimated from watershed vegetation area [19, 37] and then converted to the flux entering the drainage network using a transport coefficient that accounted for soil and riparian retention [37]. Potential additional carbon mineralization from each source was calculated as follows:

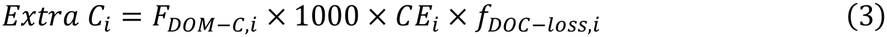

where F_DOM-C,i_ is the annual DOM-C input from source i (kt C yr^-1^); CE_i_ is the source-specific CE coefficient at the common reference addition level of 16 μM C, expressed as a decimal; f_DOC-loss,i_ is the fraction of DOC removed during the 32-d incubation for the corresponding treatment; and 1000 converts kt to t. CO_2_ equivalents were calculated by multiplying carbon mass by 44/12. Parameter sources, calculations, and uncertainty ranges are provided in Supplementary Method S4 and Tables S6 and S7.

## 3. Results

### 3.1 Source-dependent DOC degradation dynamics

During the 32-d dark incubation, the proportion of DOC remaining differed among DOM sources (Figs. 1a, S2). DOC remaining declined over time in both the lake-water control and all four source-DOM endmember incubations, although the magnitude of the decline varied among sources. By day 32, 9.9% of the initial DOC remained in the lake-water control. The corresponding proportions in the aquatic plant- and rainwater-derived DOM incubations were 5.2% and 6.0%, respectively, indicating greater DOC removal than in the control. Terrestrial-litter-derived DOM had a final DOC-remaining proportion of 8.0%, whereas sewage-derived DOM had the highest proportion remaining (11.8%) and thus the lowest DOC removal during the incubation.

**Fig. 1.**
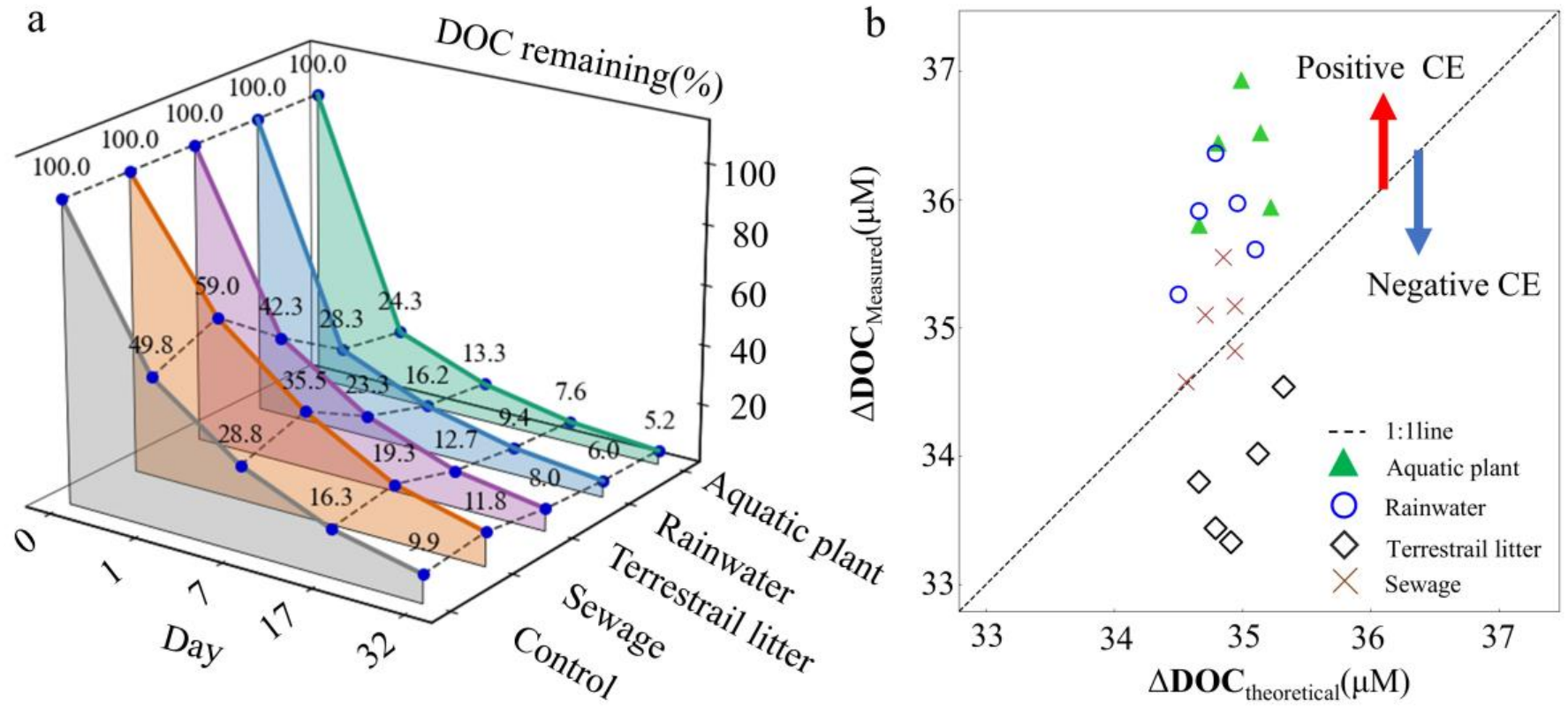
Degradation dynamics of DOM from different sources and their effects on background DOC removal. (a) Changes in the proportion of DOC remaining during the 32-d dark incubation of the lake-water control and the four source-DOM endmembers. DOC remaining is expressed as a percentage of the day-0 DOC concentration and is used to compare the degradability of background DOM with that of DOM from different sources. (b) Relationship between measured DOC removal (ΔDOC_measured_) in mixed systems and theoretical DOC removal (ΔDOC_theoretical_) under the no-interaction assumption. The dashed line is the 1:1 line. Points above the line indicate greater removal than theoretically expected (positive CE), whereas points below the line indicate lower removal than expected (negative CE). Colors and symbols denote DOM sources.

A comparison of measured DOC removal in the mixed systems with theoretical removal calculated from the lake-water and source-DOM endmembers revealed clear source-dependent deviations (Fig. 1b). Most observations for aquatic plant- and rainwater-derived DOM fell above the 1:1 line, indicating that DOC removal in the mixtures exceeded the no-interaction expectation. Terrestrial-litter-derived DOM generally remained close to the 1:1 line, with slight positive deviations at some addition levels. Sewage-derived DOM was mostly below or near the 1:1 line, indicating that DOC removal after mixing did not exceed the theoretical expectation. Thus, background DOC removal was not determined simply by the amount of exogenous DOC added but instead showed source-specific enhancement or suppression.

### 3.2 Source-dependent DOM property dynamics

The four exogenous DOM sources differed significantly in their absorbance and fluorescence properties (Fig. 2; Table S3). SUVA_254_ was relatively high for aquatic plant- and rainwater-derived DOM (3.01 ± 0.18 and 2.95 ± 0.25 L mg^-1^ m^-1^, respectively) and lowest for sewage-derived DOM (2.31 ± 0.40 L mg^-1^ m^-1^; *p* < 0.01). HIX was highest in aquatic plant-derived DOM, indicating the greatest degree of humification. In contrast, sewage-derived DOM generally had higher S_R_, BIX, β:α, and FI values, consistent with lower average molecular weight and stronger fresh autochthonous and microbial signatures. Components C1 and C3 also differed significantly among sources, further demonstrating divergence in fluorescent-component composition. Overall, aquatic plant- and rainwater-derived DOM showed stronger aromatic and humified characteristics, sewage-derived DOM showed stronger fresh autochthonous, microbial, and low-molecular-weight characteristics, and terrestrial-litter-derived DOM was intermediate.

**Fig. 2.**
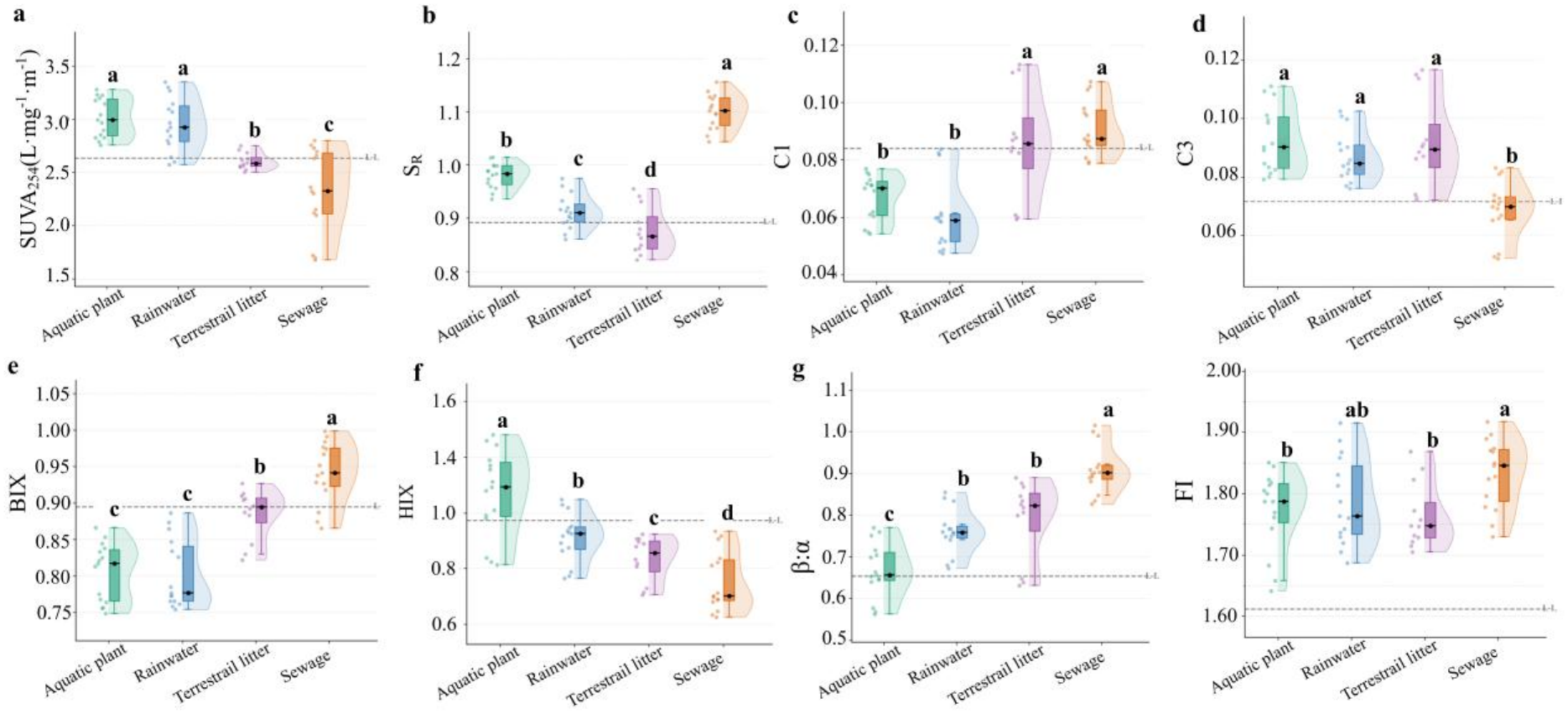
Differences in optical properties among DOM sources. Distributions of (a) SUVA_254_, (b) SR, (c) C1, (d) C3, (e) BIX, (f) HIX, (g) β:α, and (h) FI for aquatic plant-, rainwater-, terrestrial-litter-, and sewage-derived DOM. SUVA_254_ characterizes DOM aromaticity. SR is the spectral slope ratio and is related to average DOM molecular weight, with higher SR generally indicating lower average molecular weight. C1 and C3 are fluorescent components identified by PARAFAC. BIX and β:α indicate the relative contribution of fresh autochthonous DOM, HIX represents the degree of humification, and FI reflects DOM source, with higher values generally indicating stronger microbial or autochthonous contributions. Points are independent replicates; half-violin plots show probability densities; box plots show medians and interquartile ranges; and black squares show means. The gray dashed line indicates the corresponding value for background DOM in the recipient lake water. Different lowercase letters indicate significant differences among the four exogenous DOM sources (*p* < 0.05).

### 3.3 Factors influencing CE

Water-chemistry conditions at the end of the incubation differed among source treatments (Table S2). pH ranged from 7.71 to 7.92 and differed significantly among treatments (*p* < 0.001). DO ranged from 2.98 to 4.07 mg L^-1^, with the lowest value in the aquatic plant-derived DOM treatment and the highest in the sewage-derived DOM treatment (*p* < 0.01). COD_Mn_, TP, PO ^3-^-P, TN, NH_4_^+^-N, NO_2_^-^-N, and NO_3_^-^-N also differed significantly among treatments (*p* < 0.01 or *p* < 0.001). CO_2_ and N_2_O differed significantly among treatments, whereas CH_4_ did not (*p* = 0.604).

CE followed distinct source-specific trajectories across the addition gradient (Fig. 3a). For aquatic plant-, rainwater-, and terrestrial-litter-derived DOM, CE first increased and then declined as the addition level increased. The response amplitude was greatest for aquatic plant-derived DOM (R^2^ = 0.779, *p* < 0.001), followed by rainwater-derived DOM (R^2^ = 0.634, *p* < 0.01). Although weaker overall, the terrestrial-litter-derived response also showed a significant trend (R^2^ = 0.956, *p* < 0.001). Sewage-derived DOM produced negative CEs throughout the gradient, with the strongest negative response at an intermediate addition level (R^2^ = 0.559, *p* < 0.01). At 16 μM C, CE values for aquatic plant-, rainwater-, terrestrial-litter-, and sewage-derived DOM were approximately +4.4%, +3.4%, +1.5%, and −3.9%, respectively, demonstrating clear source-dependent differences in both direction and magnitude.

**Fig. 3.**
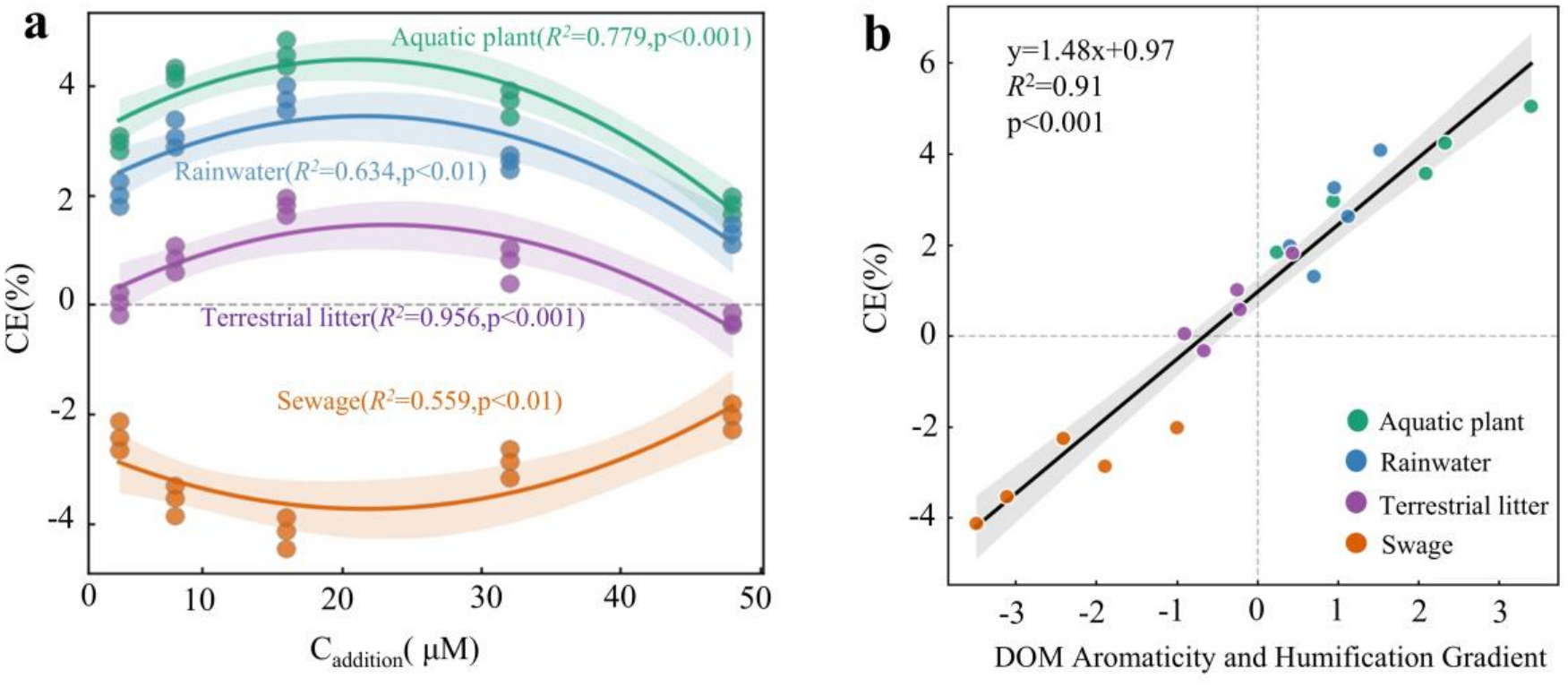
Responses of CE to addition level and its relationship with the DOM aromaticity-humification gradient. (a) Source-specific responses of the co-metabolic effect (CE) to exogenous DOM addition level. Colored fitted curves and shaded regions indicate source-specific fitted trends and their 95% confidence intervals. (b) Linear relationship between CE and the composite DOM aromaticity-humification gradient. DOM aromaticity and humification were closely related to CE direction and magnitude (R^2^ = 0.91, *p* < 0.001).

PCA further showed clear separation of DOM sources based on optical properties (Fig. S4; Table S5). PC1 and PC2 explained 51.0% and 22.0% of the variance, respectively, and source separation was jointly driven by variables including SUVA₂₅₄, C3%, HIX, BIX, and SR. After aromaticity- and humification-related indices were integrated into a composite gradient, CE was positively related to this gradient (R^2^ = 0.91, *p* < 0.001; Fig. 3b). As the gradient increased, CE shifted from negative or weakly positive values to more strongly positive values. Thus, CE direction and magnitude were closely associated with DOM optical properties: DOM with greater aromaticity and humification was more often associated with positive CE, whereas DOM with stronger fresh autochthonous, microbial, or low-molecular-weight characteristics was more often associated with negative CE.

To identify the main variables associated with CE, random forest and correlation analyses were applied to water chemistry, DOM optical properties, and gas response variables (Fig. 4). Random forest analysis indicated that variation in CE was associated with both nutrient conditions and DOM optical properties. Among water-chemistry variables, NH₄⁺-N had relatively high model importance (24.5%), whereas DO, TN, TP, NO_3_^-^-N, and pH contributed less. Among DOM optical variables, the importance values of BIX, a_350_, SUVA_254_, and HIX were 22.0%, 19.6%, 17.2%, and 13.2%, respectively.

**Fig. 4.**
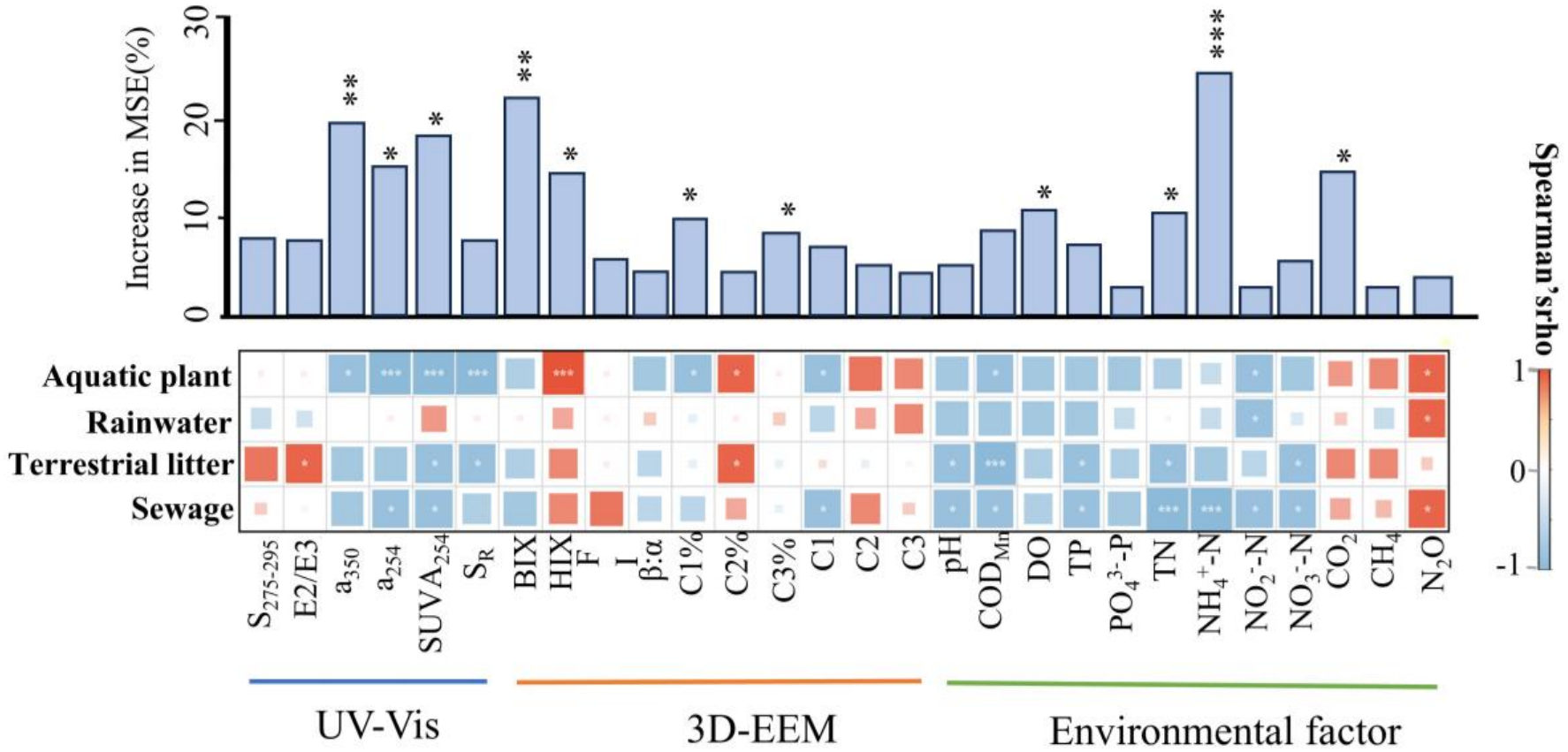
Major explanatory variables for CE and source-specific correlations. The upper bar chart shows the contribution of each variable to explaining day-32 CE in the random-forest model. Variable importance is expressed as the percentage increase in mean squared error after permutation (increase in MSE, %); larger values indicate greater explanatory contribution. The lower heat map shows Spearman correlation coefficients between CE and UV-visible absorbance indices, three-dimensional fluorescence indices, and environmental variables within each DOM source. Blue and red indicate negative and positive correlations, respectively, and color intensity indicates correlation strength. Asterisks denote statistically significant correlations. Variables on the horizontal axis are grouped into UV-visible optical indices, 3D-EEM fluorescence indices, and environmental variables; rows represent exogenous DOM sources.

Spearman correlations showed that associations between CE and environmental or DOM optical variables differed among sources (Fig. 4). Positive CE was generally more closely associated with treatments with higher SUVA_254_, HIX, and humic-like fluorescent components, whereas negative CE was more closely associated with BIX, β:α, S_R_, and stronger fresh autochthonous or microbial signatures. This pattern was consistent with the increase in CE along the aromaticity-humification gradient in Fig. 3b and further identified DOM quality as an important variable associated with differences in CE direction.

### 3.4 Potential carbon mineralization of Lake Taihu

Annual DOM-C inputs estimated from public statistics and literature-derived parameters differed markedly among sources (Fig. 5a; Table S6). Sewage- and rainwater-derived DOM were the two largest inputs, with central estimates of 5.97 and 5.32 kt C yr^-1^, respectively. Aquatic plant-derived DOM input was estimated at 2.13 kt C yr^-1^, with an uncertainty range of 0.92–3.77 kt C yr^-1^. Terrestrial-litter-derived DOM input, after correction for soil and riparian retention, had a central estimate of 0.334 kt C yr^-1^ and comparatively high uncertainty.

**Fig. 5.**
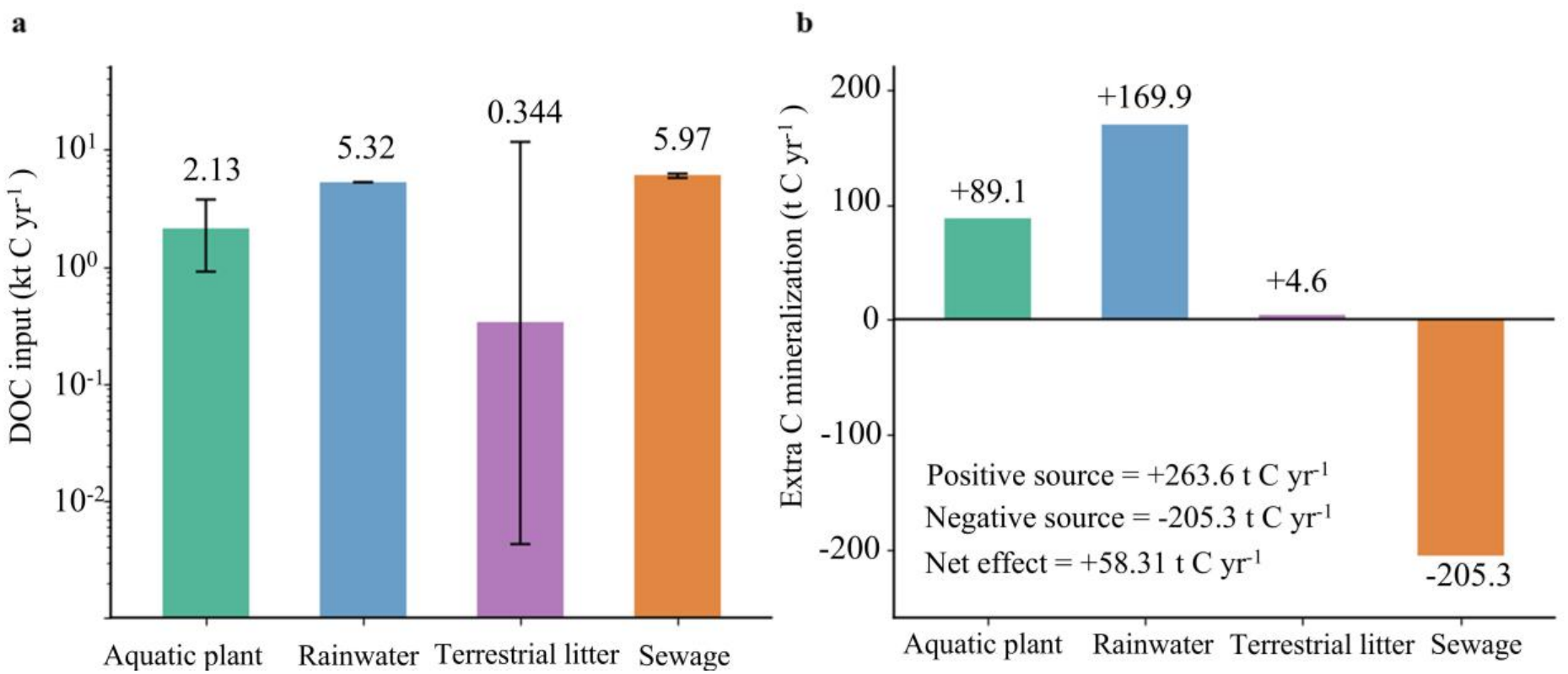
Lake Taihu-scale estimates of exogenous DOM-C inputs and the potential additional carbon mineralization they induce. (a) Annual DOM-C input fluxes from different sources to Lake Taihu, estimated from public statistics, literature-derived parameters, and mass balance. The vertical axis is logarithmic, and values are reported in kt C yr^-1^. Error bars indicate uncertainty ranges. (b) Potential source-specific additional carbon-mineralization fluxes at the lake scale, reported in t C yr^-1^. Positive values indicate additional mineralization relative to the no-interaction expectation, whereas negative values indicate lower mineralization than expected. Aquatic plant-, rainwater-, and terrestrial-litter-derived DOM made positive contributions, whereas sewage-derived DOM made a negative contribution. Net additional carbon mineralization across all four sources was +58.3 t C yr^-1^.

Combining source-specific DOM-C inputs with experimental CE values and the fraction of DOC removed during incubation yielded contrasting contributions to potential additional carbon mineralization (Fig. 5b; Table S7). Aquatic plant-, rainwater-, and terrestrial-litter-derived DOM corresponded to potential positive additional mineralization of +89.1, +169.9, and +4.6 t C yr^-1^, respectively, totaling +263.6 t C yr^-1^. Because sewage-derived DOM produced a negative CE, its estimated contribution was −205.3 t C yr^-1^. Net additional mineralization across all four sources was +58.3 t C yr^-1^, equivalent to approximately +213.8 t CO_2_ yr^-1^ based on C-to-CO_2_ stoichiometry.

## 4. Discussion

### 4.1 DOM quality regulates source-dependent CE

Exogenous DOM quality was strongly associated with both CE direction and magnitude. Aquatic plant- and rainwater-derived DOM mainly produced positive CE, terrestrial-litter-derived DOM produced a weaker positive CE, and sewage-derived DOM produced negative CE. This source-dependent pattern was broadly consistent with DOM optical properties: aquatic plant- and rainwater-derived DOM, which generated stronger positive CE, showed greater aromaticity and humification; sewage-derived DOM showed stronger fresh autochthonous and microbial signatures and lower average molecular weight; and terrestrial-litter-derived DOM was intermediate. The positive relationship between CE and the aromaticity-humification gradient, together with the relatively high random forest importance of several optical indices, indicates that source-dependent differences in CE were closely associated with DOM composition. Similar source dependence has been reported in incubations using rainwater-, plant leachate-, litter-, and sewage-derived DOM [15, 16, 17].

Positive CEs generated by aquatic plant- and rainwater-derived DOM do not imply that aromatic or humified components are intrinsically more degradable. Instead, structurally complex exogenous DOM may broaden the range of substrates available to microbial communities and promote the simultaneous processing of exogenous and resident DOM. A cross-system study of inland waters found stronger positive CE in waters containing more aromatic DOM and linked these effects to microbial taxa capable of using both labile and complex substrates [21]. Resource aromaticity can also alter bacterial succession and selection among carbon components [38], and the capacity to use high-molecular-weight DOM is unevenly distributed among bacterial communities [39]. By contrast, the relatively fresh, microbial, and low-molecular-weight components of sewage-derived DOM may be more readily used. When these substrates satisfy microbial energy requirements, the relative use of resident algal DOM may decline, resulting in preferential use or substrate substitution. The broad use of low-molecular-weight DOM by diverse aquatic bacterial communities [39] is consistent with the negative CE induced by sewage-derived DOM.

Substrate selection driven by DOM quality may also be modified by nutrient and oxygen conditions. NH₄⁺-N had relatively high model importance, and DO and nutrient concentrations differed among source treatments. Previous studies have shown that carbon and nitrogen supplies can jointly regulate aquatic DOM mineralization [40], that phosphorus availability can influence the allocation of newly added carbon between microbial assimilation and respiration [41], and that oxygen availability and exposure history can alter DOM degradation [42]. Collectively, source-specific DOM composition may first alter the availability of exogenous substrate relative to resident DOM, while nutrient and oxygen conditions may further regulate microbial allocation between the two substrate pools, thereby contributing to source-dependent CE direction and magnitude.

### 4.2 DOM addition level regulates CE magnitude

In addition to source, the amount of exogenous DOM added strongly influenced CE magnitude. For all four sources, the absolute CE response was greatest near intermediate addition levels: positive CE increased for aquatic plant-, rainwater-, and terrestrial-litter-derived DOM, while negative CE also increased in magnitude for sewage-derived DOM. At higher addition levels, positive CE from the three natural sources weakened, and the negative CE from sewage-derived DOM became less pronounced. Thus, the effect of exogenous DOM on resident carbon mineralization did not follow a simple monotonic concentration-response relationship but instead changed nonlinearly with substrate supply.

This pattern may reflect shifts in microbial resource allocation between simultaneous substrate use and substrate substitution. At low addition levels, exogenous DOM may be insufficient to relieve microbial energy or nutrient limitation and therefore may have little effect on resident DOM decomposition. Intermediate additions can provide additional available carbon without fully replacing resident substrates, thereby favoring simultaneous use of both DOM pools. As exogenous supply increases further, microorganisms may allocate more metabolic activity to the newly added substrate and reduce their relative use of resident DOM. The consistently negative CE of sewage-derived DOM is consistent with substrate substitution: its strong fresh, low-molecular-weight, and microbial signatures may make exogenous carbon a preferred substrate. Aquatic microbial experiments likewise show that low-molecular-weight carbon can be used broadly by many taxa, whereas the capacity to degrade high-molecular-weight components is restricted to particular functional groups [39].

Previous studies indicate that the effect of added organic matter on an existing carbon pool depends on substrate quality, supply, and microbial resource allocation rather than on total carbon input alone. Aquatic mixing experiments have shown that labile algal organic matter can increase microbial metabolic activity and accelerate the turnover of terrestrial litter carbon, while nutrient stoichiometry regulates the allocation of carbon between microbial assimilation and respiratory mineralization [41]. Soil incubations have also shown that the amount of labile carbon input can alter both the direction and magnitude of CE [43]. Together, these findings suggest that intermediate carbon supply may sustain the use of both exogenous and resident DOM, whereas higher supply more readily shifts microbial metabolism toward the newly added substrate and weakens CE. Addition level may therefore regulate CE magnitude by altering resource allocation between exogenous and resident DOM: intermediate supply favors simultaneous use, whereas higher supply favors preferential use of the newly added substrate.

### 4.3 Estimated carbon mineralization under source-specific DOM inputs

Combining experimental CE parameters with source-specific DOM-C inputs to Lake Taihu showed that potential additional carbon mineralization did not scale directly with input flux and may also depend on the direction of source-specific CE. Rainwater- and sewage-derived DOM were the two largest inputs, with similar annual fluxes of 5.32 and 5.97 kt C yr^-1^, respectively, yet they induced CE in opposite directions. Their estimated contributions to potential additional mineralization were therefore also opposite: approximately +169.9 t C yr^-1^ for rainwater-derived DOM and −205.3 t C yr^-1^ for sewage-derived DOM. Aquatic plant-derived DOM contributed approximately +89.1 t C yr^-1^, whereas terrestrial-litter-derived DOM contributed approximately +4.6 t C yr^-1^. The three sources with positive CE totaled approximately +263.6 t C yr^-1^, and net additional mineralization across all four sources was approximately +58.3 t C yr^-1^, equivalent to +213.8 t CO_2_ yr^-1^. Thus, potential additional carbon mineralization under multiple DOM inputs may be jointly influenced by input quantity, DOM quality, and CE direction.

### 4.4 Environmental implications

These results provide a source-resolved perspective on carbon cycling in shallow eutrophic lakes. Lake Taihu has been reported to be a substantial CO_2_ source, with emissions influenced by eutrophication, cyanobacterial blooms, and organic-carbon transformation [44]. If lake carbon budgets aggregate exogenous DOC solely by its total quantity, they may obscure source-dependent effects on resident DOM decomposition. In this study, aquatic plant- and rainwater-derived DOM produced positive CE, whereas sewage-derived DOM produced a negative CE, showing that similar input fluxes can make contrasting potential contributions to carbon mineralization. The negative contribution of sewage-derived DOM further suggests that, although anthropogenic dissolved carbon can represent an important external input [34], its carbon-cycle effect also depends on substrate properties, nutrient context, and the sequence in which microorganisms use exogenous and resident DOM [45]. Assessments of anthropogenic DOM could therefore distinguish whether it enhances, suppresses, or substitutes for decomposition of the resident carbon pool rather than considering organic-carbon loading alone. Both source and CE direction should be considered when estimating potential carbon mineralization in Lake Taihu to avoid masking differences among rainfall, vegetation, litter, and sewage inputs within the total DOC load.

The opposing CEs observed here support assessing source properties and input levels alongside bulk DOC loading when interpreting carbon processing in bloom-affected waters. Inflow-driven DOM delivery has been linked to CO_2_ emissions in Lake Taihu [8], and municipal effluent is an important source of dissolved carbon to receiving waters [34]. Source-resolved monitoring that combines DOC, optical indices, and nutrient conditions could therefore help interpret changes following rainfall, vegetation-derived inputs, and effluent discharge. The concentration-dependent responses also indicate that a single addition level need not represent all field exposures. Negative CE for sewage-derived DOM should not be interpreted as evidence that effluent discharge reduces overall carbon emissions or provides an environmental benefit; the observed effect concerns the experimental comparison and does not quantify the full carbon and nutrient consequences of effluent input.

However, this study has several limitations. First, CE was calculated from DOC mass balance without stable-isotope labeling. The relative contributions of exogenous-DOM mineralization, additional decomposition of resident DOM, and transformation into microbial biomass therefore could not be separated directly. Consequently, interpretations of substrate substitution and simultaneous use rely primarily on CE direction and DOM properties. Second, absorbance and fluorescence indices are proxies for DOM quality. Although random forest and correlation analyses identify statistical associations between variables such as NH_4_^+^-N or DO and CE, they do not establish specific molecular compositions, enzymatic processes, or independent causal mechanisms. Third, the dark microcosm incubation primarily represents microbially driven DOM decomposition and excludes photodegradation, sediment-water exchange, hydrodynamic transport, and seasonal bloom dynamics. Finally, the lake-scale estimates rely on experimental parameters from a bloom-accumulation zone, public statistics, and literature-derived coefficients. These estimates should therefore be interpreted as comparisons of potential source-specific contributions rather than as direct observations of whole-lake CO_2_ flux.

## 5. Conclusions

Using a common recipient water and a matched carbon-addition gradient, this study compared CE induced by DOM from four environmental sources. Aquatic plant- and rainwater-derived DOM produced relatively strong positive CE, terrestrial-litter-derived DOM produced a weaker positive CE, and sewage-derived DOM consistently produced a negative CE. For all sources, the absolute response was greatest at intermediate addition levels, demonstrating that both DOM source and input level regulate resident-DOM decomposition. More aromatic and humified DOM was associated with stronger positive CE, potentially favoring the simultaneous utilization of exogenous and resident DOM, whereas fresh, low-molecular-weight DOM was associated with negative CE, suggesting a preference for substrate substitution and preferential use. Additionally, nutrient and oxygen conditions may further modify these processes. Lake-scale estimates indicated that positive and negative source-specific CE can offset each other, resulting in a potential net additional mineralization of approximately +58.3 t C yr^-1^ (+213.8 t CO_2_ yr^-1^) across the four sources. Therefore, lake carbon-cycle assessments should account for DOM source, quality, and loading intensity.

## CRediT authorship contribution statement

Jiaxing Tian: Conceptualization, Methodology, Investigation, Formal analysis, Writing – original draft. Jingmin Wang: Investigation. Yujia Chen: Investigation, Formal analysis. Wenxin Wei: Investigation, Formal analysis. Xudong Dai: Investigation. Ziqian Gong: Investigation. Qiuyuan Ren: Investigation. Chao Wu: Conceptualization. Yaoqing Peng: Methodology, Writing – review & editing. Jicheng Zhong: Methodology. Fei Yang: Methodology. Xiaoguang Xu: Conceptualization, Writing – review & editing. Jianjun Wang: Methodology, Writing – original draft. All authors reviewed and approved the final manuscript.

## Data availability

Data will be made available on request.

## Acknowledgements

This study was supported by State Key Laboratory of Lake and Watershed Science for Water Security (2024SKL009), the Open Foundation of Key Laboratory of Taihu Basin Water Resources Management, Ministry of Water Resources (Yk924004), the National Natural Science Foundation of China (42577283 and 42225708), and the Natural Science Foundation of Jiangsu Province (BK20231284).

