## Supplementary Information for "Multi-source DOM inputs dictate net extra carbon mineralization in eutrophic lakes via non-linear co-metabolism effects"

**Text S1. Measurement and analysis of DOM optical properties**

Measurements of CDOM were conducted in room temperature (20 ± 2 °C) and Milli-Q water was used in the reference cell. The DOM absorption spectra at wavelengths of 200−800 nm were recorded using 1 cm path length quartz cuvettes. EEMs were measured with a xenon excitation source using a 1 cm quartz cell. The excitation wavelength was 200−450 nm at an increment of 5 nm, and emission wavelengths varied from 250 to 500 nm with 1 nm intervals. Excitation and emission silts were both set to a 5 nm bandpass. The scan speed was 6000 nm·min−1. Data was standardized and normalized to Raman Units (R.U.) based on the measurements of Raman peak at 350 nm. Data collected at excitation wavelengths below 250 nm were omitted before further analysis due to deteriorating signal-to-noise ratios. The UV-vis absorption parameters were calculated in R (R Core Team, 2022) using the package stadom (Pucher et al., 2019). Fluorescent components were characterized and identified using PARAFAC modeling and matched with previous studies using OpenFluor database (https://openfluor.lablicate.com/) (Murphy et al., 2013). To correct and calibrate the fluorescence spectra before the analysis, blank samples of deionized water was included during each analysis and Raman-normalized blanks were subtracted to remove the Raman scattering signal. Inner-filter correction was also applied. Rayleigh scatter bands were also trimmed. All valid EEMs obtained from the incubation experiment were included in the dataset modeled with PARAFAC. The EEMs data array were consisted of 41 excitation wavelengths and 251 emission wavelengths and were split into two random halves, the PARAFAC algorithm was then applied stepwise to both data arrays for three- to six- components. Ten iterations and a convergence criterion of $10^{-8}$ were used for each model. A three-component model was well-validated using half-split validation and random initialization. PARAFAC component intensities were normalized to the sum of component fluorescence intensities and expressed as their relative contribution to the total fluorescence. PARAFAC modeling was performed in MATLAB R2021a using the drEEM Toolbox (ver 0.6.0) (https://dreem.openfluor.org/) (Murphy et al., 2013).

**Text S2. Data quality**

The pH and dissolved oxygen (DO) probes of the multiparameter water quality instrument were calibrated using standard solutions prior to measurement: pH buffers of 4.01, 7.00, and 9.18 at 25 °C, and a 100% saturation point in water-saturated air for DO. Replicate measurements indicated accuracies of ± 0.05 pH units and ± 0.1 mg $L^{-1}$ for DO. Total nitrogen (TN) and total phosphorus (TP) were analyzed in replicates with an uncertainty of less than 3%. Dissolved organic carbon (DOC) was determined automatically two or three times per sample, maintaining a coefficient of variance of less than 5%. Greenhouse gas concentrations ($CO_2$, $CH_4$, and $N_2O$) were measured using a gas chromatograph (GC). The GC was calibrated using standard gas mixtures before each batch of analysis, and analytical precision was ensured to be within ± 3% through replicate injections. (APHA et al., 2023).

**Text S3. Concentration-response modeling, PCA, correlation, and random forest analysis**

To characterize changes in CE along the ordered addition gradient, polynomial regressions containing linear and quadratic concentration terms were fitted separately for each DOM source. Model residuals and goodness of fit were used to evaluate the relationships. These regressions were used to describe response trends within the experimental concentration range. Principal component analysis (PCA) was conducted using a sample-by-variable input matrix to explore the data structure. Prior to the analysis, continuous variables were screened for missing values, outliers, and dimensional heterogeneity, followed by appropriate centering and standardization (Z-score normalization). The PCA results, including eigenvalues, individual and cumulative explained variances, loading matrices, sample scores, variable retention criteria, and the construction methods for grouping ellipses, are detailed in the Supplementary Information (Jolliffe and Cadima, 2016).

Associations of CE with water chemistry, DOM optical indices, and gas variables were assessed using Spearman rank correlations. A Random Forest (RF) model was constructed to identify the principal variables explaining day-32 CE. Candidate predictors included added-DOC concentration, DOM absorbance and fluorescence indices ($a_{254}$, $a_{350}$, $SUVA_{254}$, $E_2/E_3$, $S_{275–295}$, $S_R$, FI, HIX, BIX), PARAFAC fluorescent components, pH, DO, and major nutrient indicators. Cumulative gas production was treated as a carbon-mineralization response parallel to CE and was therefore excluded from the predictor set. Variable importance was evaluated using permutation importance. Model robustness and predictive performance were evaluated using cross-validation $R^2$, root mean square error (RMSE), and model stability tests (Breiman, 2001).

### Text S4. Estimation of Lake-Scale Multi-Source DOM-C Inputs and Additional Carbon Mineralization

#### 1. Overall estimation framework

To quantify the potential role of exogenous dissolved organic carbon (DOM-C) inputs in lake-scale carbon cycling in Lake Taihu, we constructed a bottom-up, source-resolved mass-balance framework. Annual DOM-C input fluxes ($F_{DOM-C}$, kt C $yr^{-1}$) were estimated independently for four major source categories: rainwater, municipal sewage effluent, living aquatic macrophytes, and terrestrial litter. The source-specific input fluxes were then coupled with the co-metabolic effect (CE) and DOC loss fraction ($f_{DOC-loss}$) derived from the microcosm incubations to estimate potential additional carbon mineralization (Extra C) at the Lake Taihu scale (Bianchi, 2011; Cui et al., 2023; Jiang and Li, 2024).

Because no isotope tracer was used, the lake-scale calculation should be interpreted as a literature- and statistics-constrained first-order estimate of potential CE-associated carbon-mineralization consequences rather than a direct measurement of whole-lake $CO_2$ emissions. All input terms are expressed as carbon mass, and the direction of Extra C follows the sign of CE: positive values indicate additional mineralization relative to the no-interaction expectation, whereas negative values indicate an apparent offset or inhibition.

**2. Calculation methods for annual DOM-C input fluxes ($F_{DOM\text{-}C}$)**

2.1 Atmospheric deposition (rainwater DOM)

The DOC load directly entering the lake surface via rainfall was calculated from lake surface area, annual precipitation, and the DOC concentration of rainwater. To avoid unit ambiguity, $A_{lake}$ was used in $m^2$ and P was used in m:

$$F_{rain} = \frac{A_{lake} \times P \times 10^3 \times C_{rain,DOC}}{10^{12}} \qquad (S4-1)$$

where $F_{rain}$ is the rainwater DOM-C input (kt C $yr^{-1}$), $A_{lake}$ is the surface area of Lake Taihu ($2.33811 \times 10^9$ $m^2$, equivalent to 2338.11 $km^2$), P is the multi-year mean annual precipitation in the Taihu Basin (1.177 m $yr^{-1}$), and $C_{rainDOC}$ is the DOC concentration of rainwater. Lake area and precipitation were taken from Taihu-specific published data (Cai et al., 2022), and $C_{rainDOC}$ was set to 161 μM C, equivalent to 1.932 mg C $L^{-1}$, based on the global continental precipitation median reported by Willey et al. (2000). The resulting central rainwater DOM-C input was 5.32 kt C $yr^{-1}$.

2.2 Anthropogenic sewage discharge (sewage DOM)

For the anthropogenic source term, treated municipal wastewater effluent, rather than influent sewage, was used to represent the DOC load most likely to be discharged into receiving waters. The annual effluent DOM-C load was calculated as:

$$F_{sewage} = \frac{Q_{effluent} \times 10^3 \times C_{effluent,DOC}}{10^{12}} \qquad (S4-2)$$

where $F_{sewage}$ is the sewage-derived DOM-C input (kt C $yr^{-1}$), $Q_{effluent}$ is the annual volume of treated municipal wastewater effluent ($m^3$ $yr^{-1}$), and $C_{effluentDOC}$ is the DOC concentration of

WWTP effluent (mg C $L^{-1}$). We used the 2024 wastewater treatment volume of WWTPs in Wuxi City ($8.47 \times 10^8$ $m^3$ $yr^{-1}$) as the regional effluent volume, based on the municipal public report (Wuxi Municipal Bureau of Municipal Affairs and Landscaping, 2025). $C_{effluentDOC}$ was constrained using recently reported median effluent DOC concentrations for the Yangtze River Basin (6.69 mg C $L^{-1}$) and China as a whole (7.40 mg C $L^{-1}$) (Cao et al., 2025). This yielded a sewage DOM-C input range of 5.67–6.27 kt C $yr^{-1}$, with a central value of 5.97 kt C $yr^{-1}$.

2.3 Autochthonous release from aquatic plants (aquatic plant DOM)

DOC released by living aquatic macrophytes and their epiphytic communities was treated as an autochthonous DOM input pathway. This term represents living aquatic plants and periphyton DOC release, not the complete decomposition of dead macrophyte biomass. The flux was estimated from macrophyte distribution area, in situ DOC release rate, and effective sunshine hours during the growing season:

$$F_{aquatic\ plant} = \frac{A_{macrophyte} \times R_{DOC} \times T_{sun,growing}}{10^{12}} \qquad (S4-3)$$

where $F_{aquatic\ plant}$ is the aquatic-plant-derived DOM-C input (kt C $yr^{-1}$), $A_{macrophyte}$ is the area covered by aquatic vegetation ($m^2$), $R_{DOC}$ is the in situ DOC release rate (mg C $m^{-2}$ $h^{-1}$), and $T_{sun,growing}$ is the effective annual sunshine time during the growing season (h $yr^{-1}$). $A_{macrophyte}$ was constrained using the historical aquatic vegetation distribution in Lake Taihu (central value 341.3 $km^2$, range 187.5–485.0 $km^2$; Zhao et al., 2013). $R_{DOC}$ was set to $4.57 \pm 0.65$ mg C $m^{-2}$ $h^{-1}$, based on in situ measurements of submerged macrophyte-epiphyte DOC release (Demarty and Prairie, 2009). $T_{sun,growing}$ was set to 1250–1487 h $yr^{-1}$ for the April–November growing season. The resulting aquatic plant DOM-C input was 2.13 kt C $yr^{-1}$, with an uncertainty range of 0.92–3.77 kt C $yr^{-1}$.

2.4 Terrestrial-litter leaching (terrestrial-litter DOM)

A watershed-scale natural litter-derived DOC-C framework was utilized to estimate the potential DOC-C production from natural vegetated land within the Taihu Basin. The calculation incorporates land-cover area, litter dry-mass production, litter carbon content, DOC leaching yield, and a soil-filtered watershed-delivery correction. This stepwise separation ensures that the potential DOC generated by litter leaching is not directly misinterpreted as a measured lake input (Don and Kalbitz, 2005; Wang et al., 2019; Franklin et al., 2020). The watershed-scale potential terrestrial-litter DOC-C production was calculated as:

$$F_{terrestrial\ litter,potential} = \frac{\sum_k A_k \times M_{litter,k} \times f_{C,k} \times Y_{DOC,k}}{10^9} \qquad (S4-4)$$

where $F_{terrestrial\ litter,potential}$ is the watershed-scale potential terrestrial-litter leachate DOC-C production (kt C $yr^{-1}$), $A_k$ is the area of litter-producing land-cover class k ($m^2$), $M_{litter,k}$ is annual litter dry mass per unit area (g dry mass $m^{-2}$ $yr^{-1}$), $f_{C,k}$ is the carbon fraction of litter dry mass (g C $g^{-1}$ dry mass), and $Y_{DOC,k}$ is the fraction of litter carbon released as DOC during aqueous leaching (g DOC-C $g^{-1}$ litter-C). The divisor $10^9$ converts g C $yr^{-1}$ to kt C $yr^{-1}$.

For the main calculation, natural litter-producing land was divided into forest and non-forest natural vegetation/wetland classes. The 2020 Taihu Basin land-cover statistics were used to constrain $A_{forest}$ (5,580 $km^2$) and $A_{shrub\text{-}grass\text{-}wetland}$ (680 $km^2$), while cropland was excluded to avoid mixing terrestrial litter with managed crop residues. Forest litter dry-mass production was constrained at 400–700 g dry mass $m^{-2}$ $yr^{-1}$ from subtropical forest litterfall syntheses and regional forest studies. The non-forest natural vegetation/wetland litter term was constrained using the measured litter dry-mass range in Taihu lakeside forest–grass strips (20.6–119.2 g dry mass $m^{-2}$). Litter dry mass was converted to carbon using the IPCC default biomass carbon fraction of 0.47 g C $g^{-1}$ dry mass, and DOC leaching yield was set to 0.010–0.062 g DOC-C $g^{-1}$ litter-C based on fresh foliar-litter leaching experiments.

Using the lower, central and upper parameter combinations gave the following watershed potential production estimates: 10.56, 52.73 and 116.18 kt C $yr^{-1}$, respectively. These values represent potential DOC-C production from litter leaching within the watershed and should not be treated as direct lake input.

Because most litter-derived DOC can be retained or transformed by soil and riparian interfaces before it reaches surface waters, a soil-filtered watershed-exportable flux was calculated as:

$$F_{terrestrial\ litter,watershed-delivered} = F_{terrestrial\ litter,potential} \times D_{delivery} \qquad (S4-5)$$

where $D_{delivery}$ is the fraction of litter-derived DOC remaining after soil/riparian retention and potentially exportable to the drainage network. Published soil-column and rainfall-simulation studies indicate strong retention of litter-derived DOC; therefore, $D_{delivery}$ was bounded at 0.0004–0.10. Because this range spans orders of magnitude, the central plotting value was calculated with the geometric midpoint of the bounded delivery range ($D_{delivery}$ = 0.00632), rather than the arithmetic midpoint. This produced a soil-filtered watershed-delivered terrestrial-litter DOM-C flux of 0.00422–11.62 kt C $yr^{-1}$, with a central value of 0.334 kt C $yr^{-1}$. The delivered flux, not the potential production term, was used for lake-scale CE and $CO_2$-equivalent coupling. (Don and Kalbitz, 2005; Wang et al., 2019; Franklin et al., 2020)

**3. Upscaling of co-metabolic effects and conversion to additional carbon mineralization**

To estimate the potential carbon consequence of each exogenous DOM pathway after entering the water body, source-specific annual DOM-C inputs were coupled with the experimentally derived CE and DOC loss fractions. The source-specific additional carbon mineralization was calculated as: (Bianchi, 2011; Cui et al., 2023; Jiang and Li, 2024)

$$Extra\ C_i = F_{DOM-C,i} \times 1000 \times CE_i \times f_{DOC-loss,i} \qquad (S4-6)$$

where Extra $C_i$ is the potential additional carbon mineralization associated with source i (t C $yr^{-1}$), $F_{DOM\text{-}C,i}$ is the annual source-specific DOM-C input (kt C $yr^{-1}$), $CE_i$ is the source-specific CE coefficient at the common reference addition level of 16 μM C, expressed as a decimal, and $f_{DOC\text{-}loss,i}$ is the fraction of DOC removed during the 32-d incubation for the corresponding source treatment.

For comparison with greenhouse-gas accounting, carbon mass was converted to $CO_2$ mass using the molecular-weight ratio 44/12:

$$Extra\ CO_{2,i} = Extra\ C_i \times \frac{44}{12} \qquad (S4-7)$$

Using the central DOM-C inputs and the incubation-derived parameters used in the current working dataset, aquatic plant DOM generated +89.1 t C $yr^{-1}$ (+326.5 t $CO_2$ $yr^{-1}$), rainwater DOM generated +169.9 t C $yr^{-1}$ (+623.1 t $CO_2$ $yr^{-1}$), and terrestrial litter DOM generated +4.57 t C $yr^{-1}$ (+16.77 t $CO_2$ $yr^{-1}$) when the soil-filtered watershed-delivered terrestrial-litter DOM-C flux was used. In contrast, sewage DOM generated a negative CE-associated contribution of −205.3 t C $yr^{-1}$ (−752.6 t $CO_2$ $yr^{-1}$). The resulting net effect across the four source pathways was +58.3 t C $yr^{-1}$, equivalent to +213.8 t $CO_2$ $yr^{-1}$.

**Table S1. Definition and characteristics of the absorbance and fluorescence optical properties of DOM used in this study.**

| Spectral parameters | Definition | Significance | References |
|---|---|---|---|
| $a_{254}$ ($m^{-1}$) | Absorption coefficient. $a = \frac{2.303A}{l}$. A=absorbance at the wavelength of 254 nm and l = path length (m) | Used to quantify CDOM abundance. | (Helms et al., 2008; Hansen et al., 2016) |
| a350 ($m^{-1}$) | Absorption coefficient. $a = \frac{2.303A}{l}$. A=absorbance at the wavelength of 350 nm and l = path length (m) | Serves as a robust proxy for the abundance and concentration of chromophoric dissolved organic matter (CDOM). | (Helms et al., 2008; Hansen et al., 2016) |
| $SUVA_{254}$ (L $mg^{-1}$ $m^{-1}$) | Specific ultraviolet absorbance at 254 nm. $SUVA_{254} = \frac{a_{254}}{\mathrm{DOC}}$. | Higher values are associated with greater aromatic content. | (Weishaar et al., 2003; Hansen et al., 2016) |
| $S_R$ | Spectral slope ratio calculated as $S_{275-295}$ divided by $S_{350-400}$ | Negatively correlated to DOM molecular weight and DOM is mainly form biological sources when $S_R > 1$ and DOM is exogenous when $S_R < 1$. | (Helms et al., 2008; Hansen et al., 2016) |
| $E_2/E_3$ | Absorption ratio $\frac{E_2}{E_3} = \frac{a_{250}}{a_{365}}$ | The decreasing ratio indicate increasing molecular size. | (Peuravuori and Pihlaja, 1997; Hansen et al., 2016) |
| $S_{275-295}$ ($nm^{-1}$) | Slope of ln[a(λ)] over 275-295 nm | It is inversely correlated with the average molecular weight and aromaticity of DOM. | (Helms et al., 2008) |
| Fluorescence index (FI) | FI = $\lambda_{Em}$ at 470 (450) nm $\lambda_{Em}$ at 520 (500) nm at $\lambda_{Ex}$ 370 nm | Used to trace CDOM composition and source, to discriminate the sources of DOM. 1.9 for aquatic and microbial sources and about 1.4 for terrestrial and soil sources. | (McKnight et al., 2001; Hansen et al., 2016) |

| Spectral parameters | Definition | Significance | References |
|---|---|---|---|
| Humification (HIX) index | HIX = ∑435→480 nm ∑300→345 nm + ∑435→480 nm at $\lambda_{Ex}$ 254 nm | Used as an indicator for the degree of humification in DOM, and higher values indicate increasing degree of humification. HIX > 6 indicates high humification with large terrestrial contribution, HIX < 4 indicates weak humification degree and being autogenous | (Ohno, 2002; Huguet et al., 2009) |
| Biological index (BIX) | BIX = $\lambda_{Em}$ at 380 nm $\lambda_{Em}$ at 430 nm at $\lambda_{Ex}$ 310 nm | Used to quantify the proportion of freshly produced substances in DOM. BIX > 1 is biogenic, BIX < 0.7 is low native component, 0.7-0.8 is moderate native component, and 0.8-1.0 is strong native component. Positively correlate with β:α | (Huguet et al., 2009; Hansen et al., 2016) |
| Freshness index (β:α) | The ratio of emission intensity at 380 nm divided by the maximum emission intensity between 420 nm and 435 nm at excitation 310 nm | An indicator of recently produced DOM, with higher values representing a higher proportion of fresh DOM | (Parlanti et al., 2000; Hansen et al., 2016) |

**Table S2. Key environmental variables after the degradation experiment. All data are expressed as mean ± standard deviation (Mean ± SD).**

| Environmental variables | Aquatic plant | Rainwater | Terrestrial litter | Sewage | Control | *p* value |
|---|---|---|---|---|---|---|
| pH | 7.71 ± 0.13 | 7.77 ± 0.12 | 7.82 ± 0.12 | 7.92 ± 0.10 | 7.86 ± 0.10 | $p < 0.001$ |
| DO | 2.98 ± 0.34 | 3.25 ± 0.17 | 3.40 ± 0.15 | 4.07 ± 0.32 | 3.63 ± 0.05 | $p < 0.01$ |
| $COD_{MN}$ | 1.91 ± 0.19 | 2.12 ± 0.20 | 2.25 ± 0.19 | 2.92 ± 0.28 | 2.58 ± 0.03 | $p < 0.01$ |
| TP | 0.19 ± 0.00 | 0.19 ± 0.01 | 0.20 ± 0.01 | 0.21 ± 0.01 | 0.20 ± 0.00 | $p < 0.001$ |
| $PO_4^-$-P | 0.020 ± 0.003 | 0.022 ± 0.003 | 0.023 ± 0.001 | 0.036 ± 0.002 | 0.028 ± 0.000 | $p < 0.001$ |
| TN | 2.88 ± 0.07 | 2.91 ± 0.06 | 2.94 ± 0.08 | 3.08 ± 0.06 | 3.00 ± 0.04 | $p < 0.001$ |
| $NH_4^+$-N | 0.095 ± 0.011 | 0.11 ± 0.01 | 0.11 ± 0.01 | 0.14 ± 0.01 | 0.12 ± 0.00 | $p < 0.001$ |
| $NO_2^-$-N | 39.17 ± 2.55 | 41.76 ± 2.50 | 43.30 ± 2.78 | 51.82 ± 3.11 | 48.23 ± 0.61 | $p < 0.01$ |
| $NO_3^-$-N | 0.083 ± 0.007 | 0.091 ± 0.006 | 0.092 ± 0.006 | 0.13 ± 0.01 | 0.12 ± 0.00 | $p < 0.001$ |
| $CO_2$ | 10594.67 ± 351.46 | 9764.35 ± 515.57 | 8291.77 ± 372.35 | 6509.13 ± 256.53 | 7267.21 ± 91.29 | $p < 0.01$ |
| $CH_4$ | 1.86 ± 0.03 | 1.86 ± 0.02 | 1.85 ± 0.02 | 1.85 ± 0.02 | 1.83 ± 0.02 | p = 0.604 |
| $N_2O$ | 0.73 ± 0.05 | 0.67 ± 0.05 | 0.59 ± 0.03 | 0.46 ± 0.03 | 0.62 ± 0.01 | $p < 0.001$ |

**Table S3. DOM changes in different groups of the degradation experiment**

| DOM | Aquatic plant | Rainwater | Terrestrial litter | Sewage | Control | *p* value |
|---|---|---|---|---|---|---|
| $a_{254}$ | 16.14 ± 2.82 | 17.78 ± 2.55 | 18.70 ± 1.96 | 21.83 ± 2.90 | 18.26 ± 0.23 | $p < 0.001$ |
| $a_{350}$ | 8.07 ± 1.01 | 8.23 ± 0.82 | 8.66 ± 0.97 | 8.62 ± 0.70 | 8.37 ± 0.11 | $p = 0.189$ |
| $SUVA_{254}$ | 3.01 ± 0.18 | 2.95 ± 0.25 | 2.62 ± 0.08 | 2.31 ± 0.40 | 2.63 ± 0.03 | $p < 0.01$ |
| $S_R$ | 0.98 ± 0.02 | 0.91 ± 0.03 | 0.88 ± 0.04 | 1.10 ± 0.03 | 0.89 ± 0.01 | $p < 0.001$ |
| $E_2/E_3$ | 4.38 ± 0.41 | 4.19 ± 0.46 | 3.97 ± 0.27 | 5.01 ± 0.44 | 4.09 ± 0.05 | $p < 0.01$ |
| $S_{275-295}$ | 0.016 ± 0.001 | 0.016 ± 0.001 | 0.015 ± 0.001 | 0.018 ± 0.001 | 0.017 ± 0.000 | $p < 0.001$ |
| FI | 1.77 ± 0.06 | 1.79 ± 0.07 | 1.76 ± 0.05 | 1.83 ± 0.06 | 1.61 ± 0.02 | $p = 0.013$ |
| HIX | 1.27 ± 0.16 | 1.11 ± 0.09 | 1.02 ± 0.08 | 0.95 ± 0.11 | 1.17 ± 0.01 | $p < 0.001$ |
| BIX | 0.80 ± 0.04 | 0.80 ± 0.05 | 0.88 ± 0.03 | 0.94 ± 0.04 | 0.89 ± 0.01 | $p < 0.01$ |
| β:α | 0.67 ± 0.07 | 0.76 ± 0.05 | 0.78 ± 0.09 | 0.91 ± 0.06 | 0.65 ± 0.01 | $p < 0.001$ |

**Table S4. Description of the three components identified by PARAFAC analysis. The maximum fluorescence ($F_{max}$) is reported in Raman units (R.U.).**

| Component | Ex/Em (nm) | openfluor comparison | descriptions and sources of DOM | references |
|---|---|---|---|---|
| C1 | 275/310 | 15 | Protein-like, tyrosine-like. Fresh, labile and autochthonous DOM | (Coble, 2007; Cory and McKnight, 2005; Fellman et al., 2010) |
| C2 | 335/440 | 17 | Humic-like. Terrestrial and highly humified DOM | (Coble, 2007; Fellman et al., 2010; Murphy et al., 2013) |
| C3 | 385/480 | 6 | Red-shifted humic-like. Strongly humified and relatively refractory DOM | (Coble, 2007; Cory and McKnight, 2005; Fellman et al., 2010) |

**Table S5. Summary of principal component analysis (PCA) results for DOM optical parameters.**

| PC axis | Eigenvalue | Explained variance (%) | Cumulative variance (%) | Dominant positive loadings |
|---|---|---|---|---|
| PC1 | 3.06 | 51.0% | 51.0% | $SUVA_{254}$, C3%, HIX |
| PC2 | 1.32 | 22.0% | 73.0% | BIX, $S_R$, $SUVA_{254}$, C3% |

**Table S6. Summary of parameters used to estimate annual source-resolved DOM-C fluxes.**

| Source | Estimation concept | Formula | Parameters | Result | Evidence and boundary |
|---|---|---|---|---|---|
| Rainwater DOM | Direct rainwater DOC deposited on Lake Taihu surface | $F_{\text{rain}} = \frac{A_{\text{lake}} \times P \times 10^3 \times C_{\text{rain,DOC}}}{10^{12}}$ | $A_{lake}$ = 2338.11 $km^2$; P = 1177 mm $yr^{-1}$; $C_{rainDOC}$ = 161 µM = 1.932 mg C $L^{-1}$ | 5.32 kt C $yr^{-1}$ | Lake Taihu area and precipitation from Cai et al. (2022); rainwater DOC concentration from Willey et al. (2000). Boundary: direct rainfall deposition onto the lake surface only. |
| Sewage DOM | WWTP effluent DOC load; influent DOC is not used as in-lake load | $F_{\text{sewage}} = \frac{Q_{\text{effluent}} \times 10^3 \times C_{\text{effluent,DOC}}}{10^{12}}$ | Q = 8.47 × $10^8$ $m^3$ $yr^{-1}$; C = 6.69-7.40 mg C $L^{-1}$ | 5.67-6.27 kt C $yr^{-1}$; central 5.97 | Wuxi WWTP treatment volume from Wuxi Municipal Bureau of Municipal Affairs and Landscaping (2025); effluent DOC medians from Cao et al. (2025). Boundary: regional treated-effluent load; apply discharge-to-Taihu routing if available. |
| Aquatic plant DOM | Living macrophyte-periphyton DOC release, not complete decay of dead biomass | $F_{\text{aquatic plant}} = \frac{A_{\text{macrophyte}} \times R_{\text{DOC}} \times T_{\text{sun,growing}}}{10^{12}}$ | A = 341.3 $km^2$ (187.5-485.0); R = 4.57 ± 0.65 mg C $m^{-2}$ $h^{-1}$; T = 1250-1487 h $yr^{-1}$ | 2.13 kt C $yr^{-1}$; range 0.92-3.77 | Aquatic vegetation area from Zhao et al. (2013); macrophyte-epiphyte DOC release rate from Demarty and Prairie (2009). Boundary: living macrophyte/periphyton release rather than complete decay of dead biomass. |

| Source | Estimation concept | Formula | Parameters | Result | Evidence and boundary |
|---|---|---|---|---|---|
| Terrestrial litter DOM | Watershed-scale natural litter-derived DOC-C production and soil-filtered delivery; cropland straw/crop residues excluded unless the endpoint is renamed. | $F_{\text{terrestrial litter,potential}} = \frac{\sum(A_k \times M_{\text{litter},k} \times f_{C,k} \times Y_{\text{DOC},k})}{10^9}$; $F_{\text{terrestrial litter,watershed-delivered}} = F_{\text{terrestrial litter,potential}} \times D_{\text{delivery}}$. | $A_{forest}$ = 5,580 km$^2$; $A_{shrub\text{-}grass\text{-}wetland}$ = 680 km$^2$; $M_{forest}$ = 400–700 g dry mass m$^{-2}$ yr$^{-1}$; $M_{herb}$ = 20.6–119.2 g dry mass m$^{-2}$ yr$^{-1}$; $f_C$ = 0.47; $Y_{DOC}$ = 0.010–0.062; $D_{delivery}$ = 0.0004–0.10. | Potential production: 10.56–116.18 kt C yr$^{-1}$; central 52.73. Soil-filtered watershed-delivered flux: 0.00422–11.62 kt C yr$^{-1}$; central 0.334. | Land-cover area from Taihu Basin 30-m land-cover data and official basin-scale report (Yang and Huang, 2021; Taihu Basin Authority of Ministry of Water Resources, 2023); forest litterfall from subtropical/regional forest litter studies (Liu et al., 2024; Zhao et al., 2022); local herbaceous litter from Liu et al. (2012); carbon fraction from IPCC (2006); DOC leaching yield from Don and Kalbitz (2005); delivery fraction from Wang et al. (2019) and Franklin et al. (2020). Boundary: use delivered flux for CE coupling; report potential production as upper process capacity. |

**Table S7. Estimates of CE-associated additional carbon mineralization and equivalent $CO_2$ emissions at the lake scale.**

| Source | DOM-C input (kt C $yr^{-1}$) | CE | DOC loss fraction | Extra C (t C $yr^{-1}$) | Extra $CO_2$ (t $CO_2$ $yr^{-1}$) |
|---|---|---|---|---|---|
| Aquatic plant | 2.135 | +0.044 | 0.948 | +89.1 | +326.5 |
| Rainwater | 5.317 | +0.034 | 0.940 | +169.9 | +623.1 |
| Terrestrial litter | 0.334 | +0.015 | 0.914 | +4.57 | +16.77 |
| Sewage | 5.967 | −0.039 | 0.882 | −205.3 | −752.6 |
| Net | — | — | — | +58.3 | +213.8 |

Note. Extra C was calculated using the central source-specific DOM-C input, source-specific CE, and source-specific DOC loss fraction (Extra $C_i = F_{\mathrm{DOM-C},i} \times 1000 \times CE_i \times f_{\mathrm{DOC-loss},i}$). Extra $CO_2$ equivalents were calculated using the molecular-weight ratio 44/12. Positive values indicate additional mineralization relative to the no-interaction expectation, whereas negative values indicate apparent offset or inhibition.

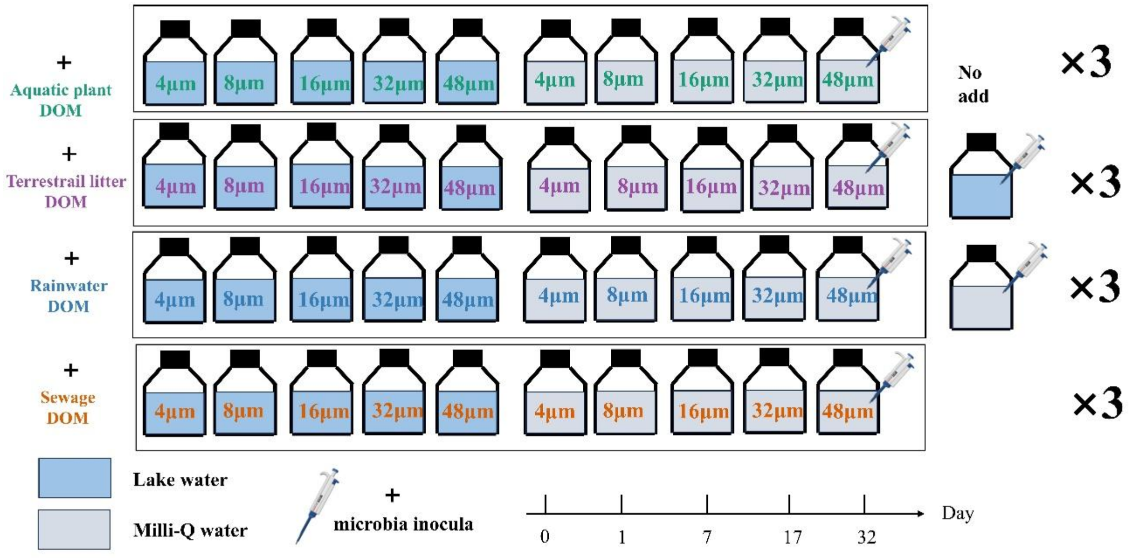


**Figure S1. Schematic diagram of the microcosm incubation experimental design.** Four exogenous dissolved organic matter (DOM) sources (aquatic plant, terrestrial litter, rainwater, and sewage) were spiked into natural lake water and Milli-Q water matrices at five dissolved organic carbon (DOC) concentration gradients (4, 8, 16, 32, and 48 μM). Standardized microbial inocula were introduced into all incubation bottles (indicated by the syringe icon). Two "No add" groups served as baseline controls for the lake water (native DOM background) and Milli-Q water matrices, respectively. All experimental treatments and controls were performed in independent triplicates (n = 3). The microcosms were incubated in the dark for 32 days, with temporal sampling conducted on days 0, 1, 7, 17, and 32 to monitor carbon mineralization and optical dynamics.

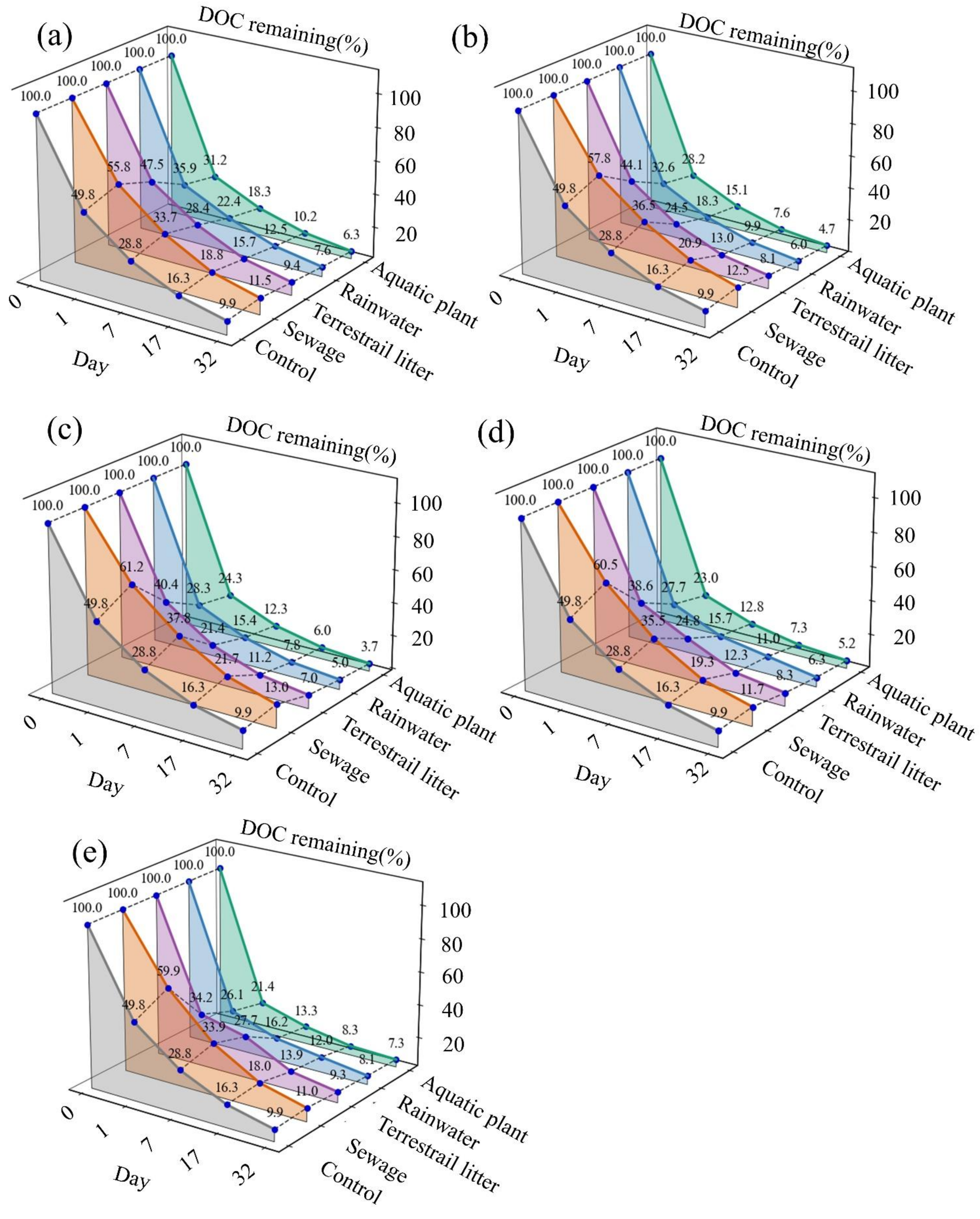


**Figure S2. Time-series of DOC remaining percentage across different DOM sources and addition concentration gradients.** Panels (a)-(e) illustrate the dynamics for five concentration gradients: (a) 4μM, (b) 8 μM, (c) 16 μM, (d) 32 μM, and (e) 48 μM. The gray line and shaded area labeled "Ctrl" denote the decomposition of native DOM in the natural lake water matrix (control group). The colored lines and areas represent the total DOC consumption dynamics following the addition of exogenous DOM from distinct sources into the lake water matrix: orange = sewage DOM, purple = terrestrial litter DOM, blue = rainwater DOM, and green = aquatic plant DOM. The temporal dynamics were monitored over 32 days, with sampling conducted on days 0, 1, 7, 17, and 32.

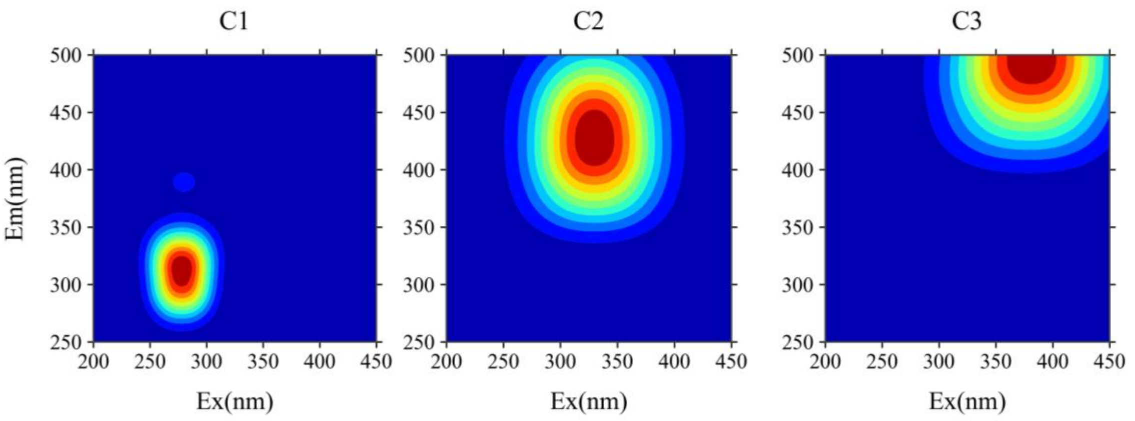


Figure S3. Three fluorescent components identified using PARAFAC analysis. The three-component model was well-validated using the split-half validation procedure. Excitation and emission peaks positions of the EEMs of the independent components are indicated alongside descriptions in Table S5.

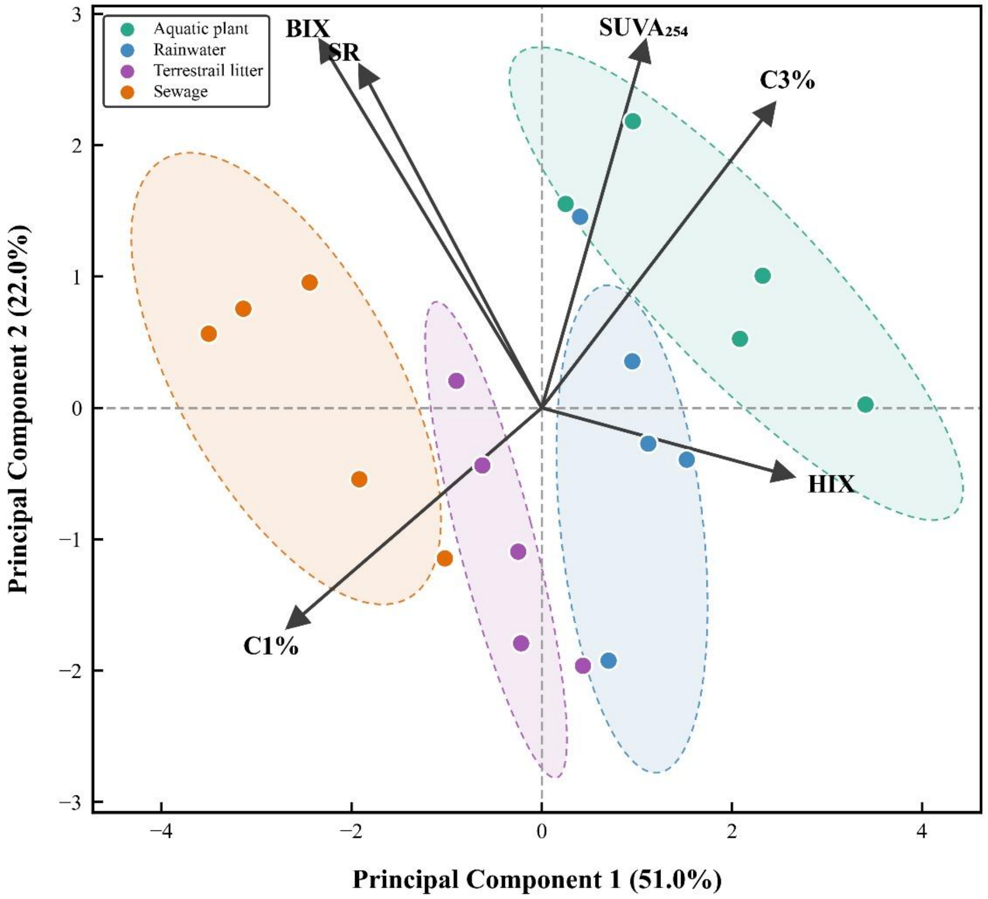


**Figure S4. Principal component analysis (PCA) of DOM optical properties across four exogenous sources.** The biplot illustrates the distribution of DOM samples (points) and the loadings of optical indices (arrows) along the first two principal components (PC1: 51.0%; PC2: 22.0%). Shaded ellipses represent the 95% confidence intervals for each source group (n = 3). Source separation was associated with loadings of $SUVA_{254}$, C3%, HIX, BIX, and $S_R$.

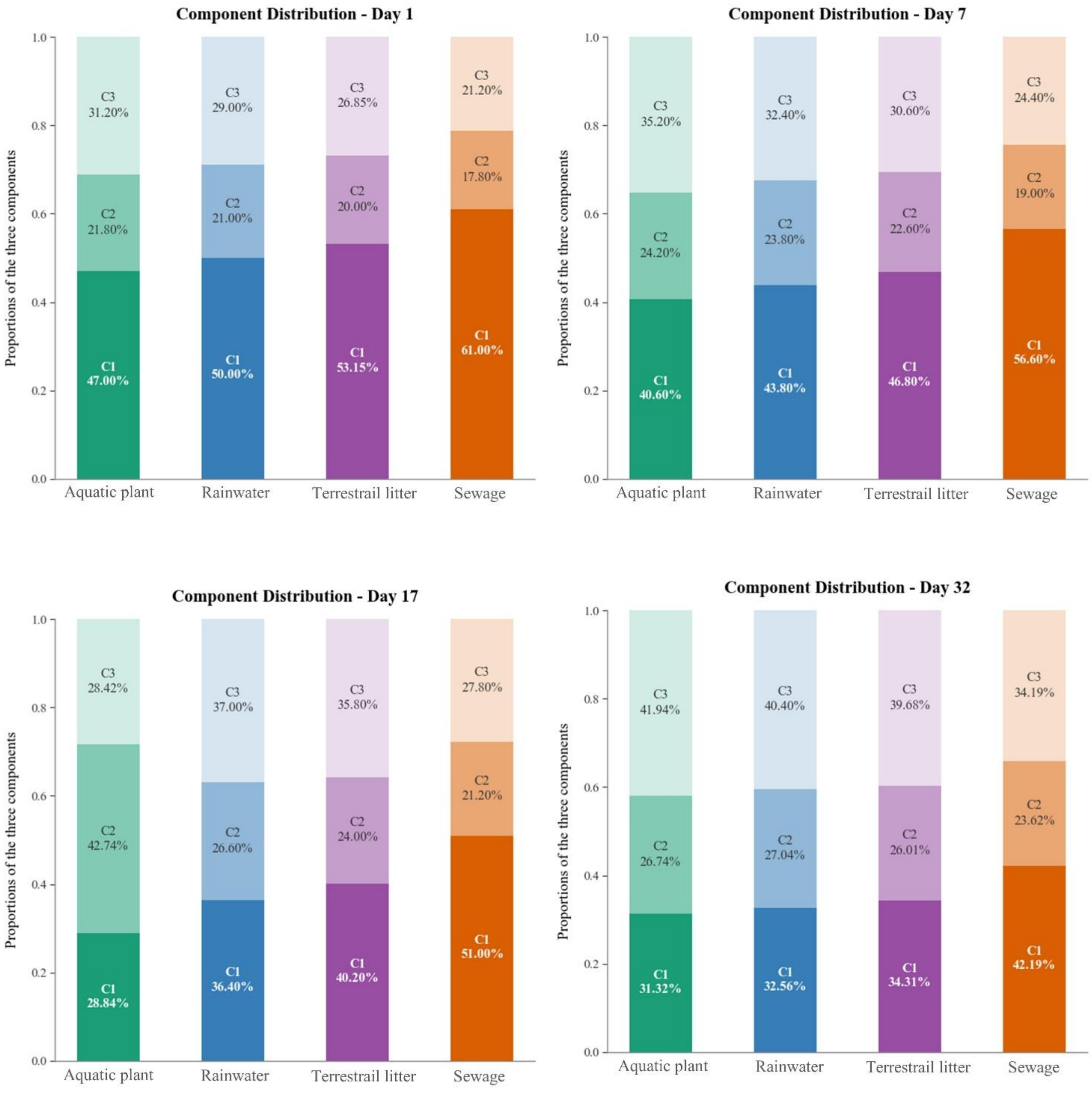


**Figure S5. The proportion of three fluorescent components identified using the PARAFAC modelling.** C1 was protein-like, whereas C2 and C3 were humic-like components.
